# Tuning of cytokine signaling through imbalanced abundances of receptors and kinases

**DOI:** 10.1101/2022.10.07.511173

**Authors:** Léa Sta, Guillaume Voisinne, Jesse Cotari, Michael F. Adamer, Carmen Molina-París, Grégoire Altan-Bonnet

## Abstract

Cells use surface receptors to detect extra-cellular ligands (*e*.*g*., growth factors or cytokines) and engender signaling cascades. We explore the effect of cell-to-cell variability in receptor subunit copy number for cells responding to *γc* cytokines (*e*.*g*., IL-2, IL-7 and IL-15). We find that primary T cells expressing higher levels of the common *γ*_*c*_ receptor chain have weaker responses to IL-7, both in terms of lowered STAT5 phosphorylation amplitude and higher EC_50_. A mathematical model that accounts for abundance imbalance (*e*.*g*., insufficient expression of JAK kinases compared to the number of receptors, or *γ*_*c*_ competition for other receptor subunit chains) predicts the formation of non-signalling complexes, consistent with the observed cellular behaviour. This type of built-in limit on signaling responses illustrates how phenotypic heterogeneity generates biological functional diversity.

**Significance Statement:** Cells rely on cytokines to coordinate their activation, differentiation, proliferation and survival. In particular, *γ*_*c*_ cytokines (interleukins IL-2, 4, 7, 9, 15, and 21) regulate the fate of leukocytes. The signaling cascade induced by these cytokines is relatively simple, and involves the phosphorylation of receptor-associated Janus-like kinases (JAK). Here, we explore the cell-to-cell variability of cytokine responses in primary mouse T cells, and find a paradoxical and quantitative imprint of receptor expression levels and other signaling components. For instance, high abundance of the common *γ*_*c*_ chain reduces cytokine responses (both in terms of signaling amplitudes and characteristic cytokine concentrations triggering 50% of the response). We develop mathematical models to quantify how limited abundances of signaling components (*e*.*g*., JAK or other cytokine receptor subunit chains) may explain our experimental observations. We conclude by generalizing this observation of cell-to-cell signaling variability to other ligand-receptor-kinase systems.

## Introduction

Biological systems have developed a variety of mechanisms for converting external information into cellular responses, including ion channels, pressure sensors, G-protein coupled receptors, receptor tyrosine kinases (RTKs), and receptors requiring associated kinases, such as the Janus kinase (JAK) family of cytokines (1, 2). The latter two families of receptors are similar in many regards. Both RTKs and JAK-associated receptors are activated by ligand-induced dimerization, which leads to phosphorylation of the receptor intra-cellular chains. This phosphorylation event results in the activation of downstream kinases, such as phosphatidylinositol-3-kinase (PI3K) and Ras, as well as the activation of proximal signal transducers and activators of transcription (STATs) (3).

In spite of their similarities, these two receptor families have one notable difference in the mechanisms by which they initiate signaling: while RTKs have intrinsic kinase activity, cytokine receptors co-opt JAK family kinases to gain signaling ability. Though, at a first glance, the difference between a kinase that is intrinsic to the receptor and one that is tightly bound to the receptor may seem negligible in terms of biochemistry, our prior studies have shown that cytokine receptor signaling can be significantly altered by the precise abundances of its molecular signaling components (4). Hence, we hypothesized that cytokine receptors and their co-optation of soluble kinases could elicit large non-genetic variability of ligand detection within a **1–14** population of cells (4, 5). We, thus, combined single-cell experimental measurements and mathematical modeling to explore the impact of the relative abundance of signaling components on the sensitivity and amplitude in response to extra-cellular cytokines (6, 7).

We focus on the common gamma (*γ*_*c*_) family of cytokine receptors, comprising the receptors for IL-2,4,7,9,15, and 21 (8). This family shares the eponymous *γ*_*c*_ chain, a ligand-binding chain without intrinsic kinase activity. The *γ*_*c*_ chain signals instead through the action of an associated JAK kinase, JAK3. All cytokine receptors in this family complete the signaling core of the receptor with a second chain associated with JAK1: IL-*x* (*x*=4,7,9 or 21) makes use of IL-*x*R*α*. IL-2 and IL-15 share the JAK1-associated IL-2R*β* chain, and include a third chain for specificity and enhanced sensitivity (IL-2R*α* and IL-15R*α*, respectively). Our previous work demonstrated that sharing the *γ*_*c*_ chain led to correlated cellular sensitivity in the *γ*_*c*_ family cytokines: increased abundance of the IL-2R*α* subunit chain decreased a cell’s sensitivity to IL-7 (4). Our computational models and subsequent experiments indicated that IL-2R*α* decreased sensitivity to IL-7 by sequestering the *γ*_*c*_ chain into pre-formed IL-2 receptor complexes (4).

Here we explore a conjecture derived from our initial finding of *γ*_*c*_ chain competition; namely, that increases in *γ*_*c*_ chain abundance would increase the number of fully-formed receptors and the sensitivity to the *γ*_*c*_ family cytokines. Furthermore, by analogy to the well-established role of RTK upregulation in cancer (9, 10), a natural expectation would be that *γ*_*c*_ chain upregulation induces greater maximal signaling upon ligand binding (1). This work, however, stems from follow-up experiments examining the impact of *γ*_*c*_ chain abundance on T cell responses to IL-7, which led us to the paradoxical observation that increased abundance of the *γ*_*c*_ chain, in fact, decreased both the sensitivity and the amplitude. Making use of mathematical modeling and quantitative flow cytometry data, we demonstrate that part of this counter-intuitive behavior is an intrinsic feature of a receptor requiring an associated kinase. Specifically, the composition of the IL-7 receptor signaling core, comprising two distinct subunit receptor chains and an associated JAK kinase (JAK3), induces a subtle balance between the different molecular components of the receptor-ligand system, required for optimal signaling. Furthermore, we also show that unbalanced expression (upregulation) of any one component is generally insufficient to increase the IL-7 response. To put this observation in perspective, we mathematically investigate the effect of chain copy number variability for a wider class of receptor types: heterodimeric or homodimeric receptors with intrinsic kinase activity, or requiring JAK kinases, and monomeric receptors. This analysis illustrates that the composition of the *γ*_*c*_ family of receptors provides functional diversity in the face of variability in molecular abundances, and demonstrates how the exact biochemical architecture of a receptor may have implications for the oncogenic potential of receptor over-expression.

## Results and discussion

### Effect of *γ*_*c*_ chain abundance on IL-7 response

To investigate the impact of the abundance of the *γ*_*c*_ cytokine receptor on cell signaling, we analyzed the response of single primary T cells to cytokine stimulation. We first assessed cellular responses to the IL-7 cytokine by flow cytometry using STAT5 phosphorylation. To probe cellular responses over a broad range of *γ*_*c*_ abundances, we transfected primary T cells with plasmids expressing *γ*_*c*_ or GFP. As compared to control cells, *γ*_*c*_ transfected cells showed elevated abundance of the receptor (Fig. S1A). Control and *γ*_*c*_ transfected cells with a similar *γ*_*c*_ abundance exhibited similar responses to IL-7 stimulation (Fig. S1B), suggesting that the transfection did not alter other components of the signaling machinery. Interestingly, cells with a high *γ*_*c*_ abundance showed reduced STAT5 phosphorylation levels, suggesting that this cytokine receptor could have a negative impact on IL-7R signaling.

To further characterize how IL-7R signaling is regulated by the abundance of its signaling components, we quantified simultaneously the abundances of *γ*_*c*_, IL-7R*α* and JAK3 (Fig. S1C-D). Analysis of single-cell responses under a high dose of IL-7 (10 nM) demonstrated heterogeneous degrees of STAT5 phoshorylation that correlated monotonically with the expression levels of IL-7R*α* and JAK3 (Fig. 1A). In contrast, and consistently with our previous observation, greater abundance of *γ*_*c*_ subunit chains was associated with reduced levels of phosphorylated STAT5 (pSTAT5).

**Fig. 1.**
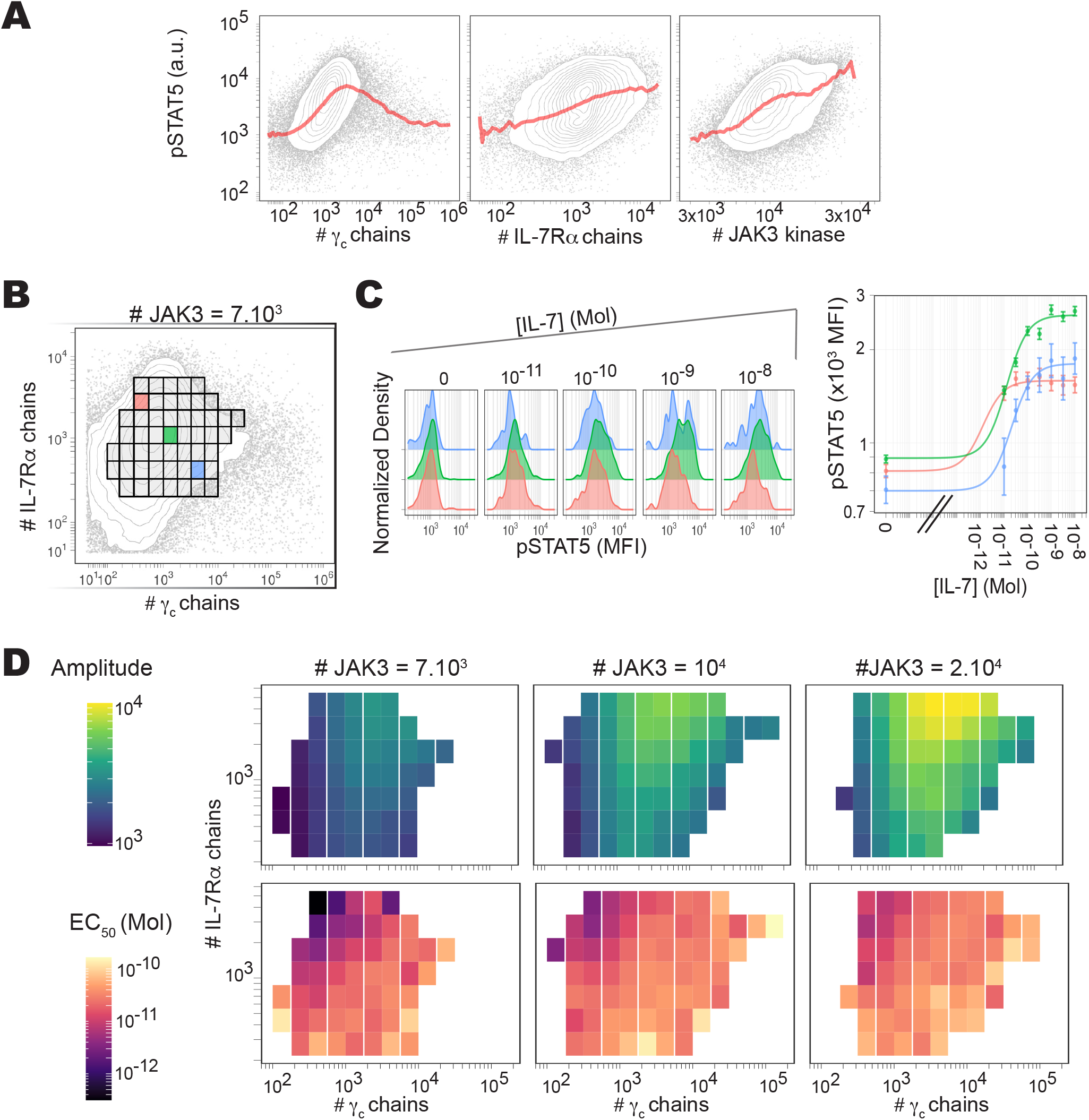
Cell-to-cell variability analysis of the pSTAT5 response to IL-7. A) Distribution of STAT5 phosphorylation (pSTAT5) in response to 10 nM of IL-7, for different abundances of IL-7R*α*, JAK3, and *γ*_*c*_ (red line represents the geometric mean of pSTAT5 for each level of signaling component). B) Definition of three emblem subpopulations based on surface abundance of IL-7R*α* and *γ*_*c*_ subunit chains. C) pSTAT5 distribution for increasing concentrations of IL-7 for the three emblem subpopulations. D) Complete maps of pSTAT5 amplitude (top) and EC_50_ (bottom) for different expression levels of the signaling components: IL-7R*α*, JAK3, and *γ*_*c*_.

To determine the relative contribution of each protein, we performed cell-to-cell variability analysis (CCVA), a method that systematically parses the effect of protein abundance on downstream responses to a stimulus (4, 11). In this case, CCVA was used to quantify the effects of *γ*_*c*_, IL-7R*α* and JAK3 abundance on T cell responses to IL-7. Cells were grouped into bins according to their abundance of *γ*_*c*_, IL-7R*α* and JAK3 (Fig. 1B). For each subpopulation, the average response to IL-7 stimulation, as measured by pSTAT5, was fitted with a Hill function and characterized in terms of its amplitude (the STAT5 phosphorylation level corresponding to the maximum IL-7 dose) and its EC_50_ (the dose of IL-7 that elicits half of the amplitude) (Fig. 1C). The output of this process is a set of maps of the relative effect of IL-7R*α, γ*_*c*_ and JAK3 abundance on both amplitude and EC_50_ (11) (Fig. 1D).

The resulting maps revealed a strikingly complex relationship between *γ*_*c*_ levels and IL-7 response. The EC_50_ map (Fig. 1D, bottom) indicated that a greater abundance of *γ*_*c*_ increased EC_50_ values. The amplitude map exposed an even more complex relationship between *γ*_*c*_ expression levels and STAT5 phosphorylation in response to IL-7 stimulation (Fig. 1D, top). A clear trend could be observed for any abundance of IL-7R*α* and JAK3, in which increasing the abundance of the *γ*_*c*_ chain first increased and then decreased the amplitude. This showed that the reduced pSTAT5 levels observed at a high IL-7 dose (10 nM) in Fig. 1A and Fig. 1C can be associated with elevated *γ*_*c*_ abundance, independently of that of IL-7R*α* and JAK3.

### A simple model of imbalance between cytokine receptor and kinase abundances (model I)

Our experimental results present a paradox: high abundance of the *γ*_*c*_ chain, an essential IL-7 receptor component, is found to reduce the responsiveness to this cytokine, reflected both in a smaller amplitude and a greater EC_50_ of the pSTAT dose-response curve. To resolve this, we considered the fact that *γ*_*c*_ does not have intrinsic kinase ability, but rather depends on the kinase activity of receptor-associated JAK3 to activate downstream signaling (2). As such, we hypothesized that increased abundance of the *γ*_*c*_ chain alone could exert a dominant negative effect via the formation of signaling-deficient receptors (or “dummy” complexes in this paper) deprived of the JAK3 kinase (Fig. 2A). Such a negative effect of over-abundance has been previously reported in the case of the CD8 co-receptor, which requires an associated downstream kinase (Lck, the lymphocyte specific protein tyrosine kinase) (12). To test this hypothesis, we modified our mathematical model of IL-7 signaling presented in (4) to account for the binding of JAK3 to the *γ*_*c*_ chain. The model, depicted in Fig. 2B and which we will refer to as model I, considers affinity constants and molecular abundances (Fig. 2C) which had been previously determined (4) making use of experimentally-validated approximations (Supplementary Table S1). Our theoretical analysis of model I, presented in (13), showed that the amplitude (the number of ligand-bound signaling receptors for a large IL-7 concentration) is equal to the total copy number of the trans-membrane chain (IL-7R*α* or *γ*_*c*_) with the lowest availability (limiting chemical reactant), modulated by a ratio that takes a value between 0 and 1. This ratio only depends on the total number of JAK3 and *γ*_*c*_ molecules and their affinity constant, *K*_1_; thus, clearly demonstrating that the abundance of JAK3 does have an influence on the amplitude, in accordance with our experimental results. Moreover, our analytic study showed that the EC_50_ value is independent of JAK3 abundance (13), in agreement with the experimental data.

**Fig. 2.**
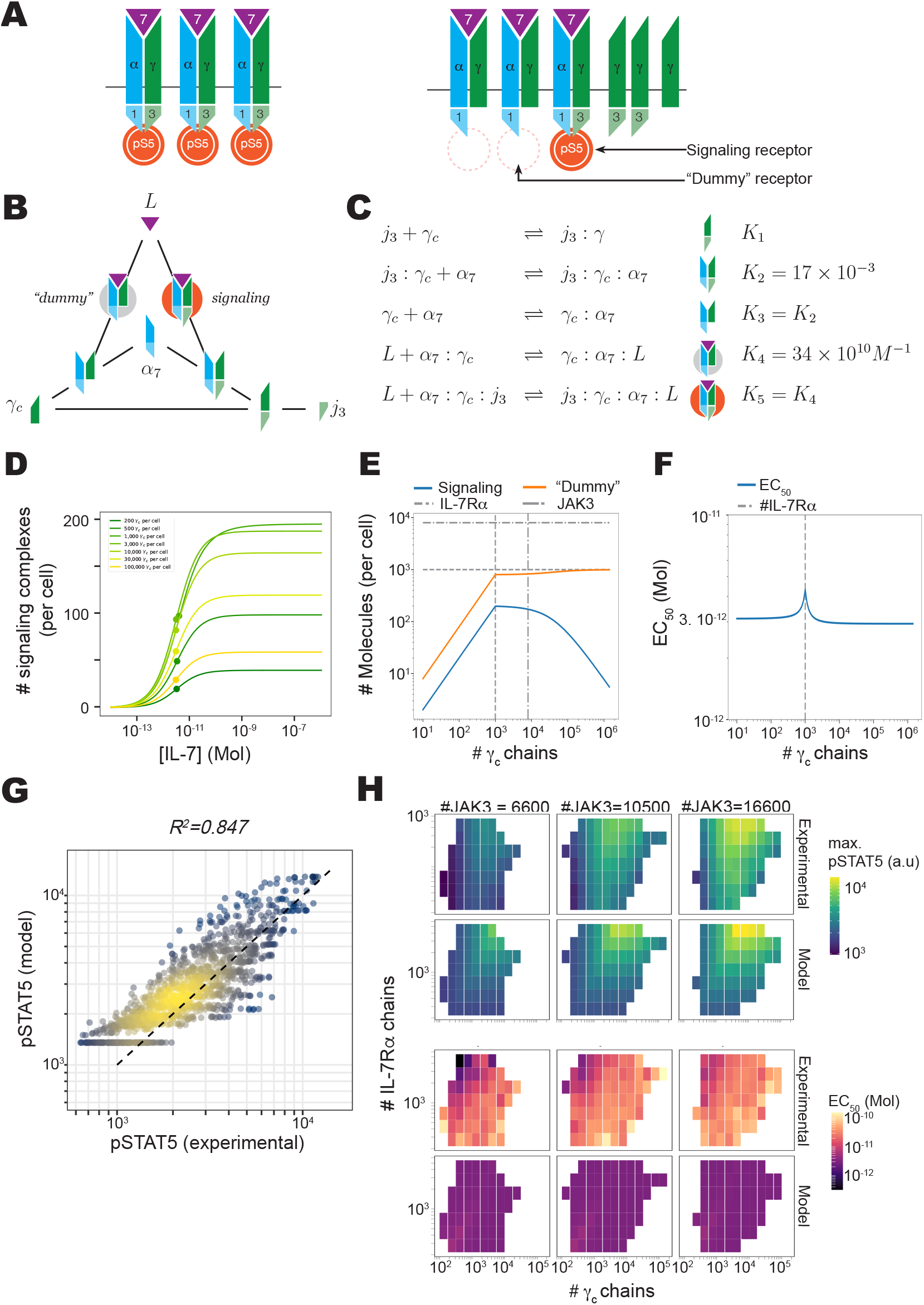
Model I of stoichiometric imbalance of JAK3 compared to *γ*_*c*_ accounts for the formation of non-signaling (or “dummy”) receptor complexes and the loss of pSTAT5 response for high copy numbers of *γ*_*c*_. A) Cartoon of the formation of the IL-7/IL-7 receptor complex. Note that some of the fully engaged receptors will fail to signal due to the absence of JAK3. B) Sketch of the biochemical reactions associated with the formation of the IL-7/IL-7R complex. C) Chemical equilibrium for the reactions depicted in B (*j*_3_: JAK3, *α*_7_: IL-7R*α, L*: IL-7). *K*_*i*_ are affinity constants for *i* = 1, …, 5. D) Modeled pSTAT5 in response to increasing doses of IL-7 (for different abundance of *γ*_*c*_ chains). E) Amplitude of responses: *i*.*e*., number of fully signaling IL-7/IL-7R (blue) or “dummy” (orange) complexes for increasing abundance of *γ*_*c*_ at high ligand concentration. F) EC_50_ for pSTAT5 in response to IL-7 for increasing abundance of *γ*_*c*_. It is essentially constant and fails to reproduce the experimental observations. G) model I is broadly accurate in predicting pSTAT5 for a wide range of IL-7 concentrations, across different cellular subpopulations. H) Model I can account for the (IL-7R*α* and *γ*_*c*_) dependence of pSTAT5 amplitude, but fails to reproduce the dependence for EC_50_ (theoretical and experimental results to be compared). Theoretical results correspond to simulations of model I with best-fit parameters corresponding to the entry “model_I” in Supplementary file Supp_table_1.

Median JAK3 abundance was measured using flow cytometry to be 8*×*10^3^ proteins per cell (see Supplementary section H). The median copy number of IL-7R*α* was reported to be 10^3^ proteins per cell (14). We use these two values as default when studying model I. We analysed model I numerically to explore how *γ*_*c*_ abundance impacts the cytokine response (Fig. 2D, E and F, Supplementary section C, and section G). As conjectured, model I demonstrated that the over-abundance of *γ*_*c*_ chains could indeed inhibit the IL-7 signaling response and thus, reduce its amplitude (Fig. 2D). Namely, both the number of signaling and non-signaling (or “dummy”) complexes grow linearly as *γ*_*c*_ chain copy number increases (Fig. 2E). When the number of *γ*_*c*_ chains equals the number of IL-7R*α* chains, the amplitude of the IL-7 response reaches its maximum. As the abundance of *γ*_*c*_ exceeds that of IL-7R*α*, the number of signaling complexes decreases and that of “dummy” complexes, in turn, increases. In our analytic study (13), we showed that, at high cytokine (IL-7) concentration, the sum of the “dummy” and signaling complexes is always equal to the total copy number of the limiting component (*γ*_*c*_ during the linear growth, and IL-7R*α* after the maximum amplitude has been reached). This implies that, once the number of *γ*_*c*_ chains exceeds that of IL-7R*α*, all IL-7R*α* chains are ligand- and *γ*_*c*_-bound to form either a signaling or a “dummy” complex. When the number of *γ*_*c*_ chains exceeds that of JAK3 molecules, the number of signaling complexes goes back to zero rather quickly (proportionally to the inverse of the total number of *γ*_*c*_ chains, as derived from the analytic expression of the amplitude in the limit when the copy number of *γ*_*c*_ tends to +*∞* (13)), and the IL-7R*α* chains are used to form “dummy” complexes, without signaling ability. This theoretical result suggests that when *γ*_*c*_ is in excess (compared to the other subunit chains of the receptor complex), it could indeed be inhibitory to the IL-7 signaling amplitude. After fitting the model to the experimental data, we found that model I could reproduce the pSTAT5 response in individual cells with accuracy (Fig. 2G).

However, if model I could account for the observed dependency of the maximal pSTAT5 on the abundance of IL-7R*α*, JAK3 and *γ*_*c*_, it could not reproduce the experimental increase of EC_50_ with increasing expression of *γ*_*c*_ chains (Fig. 2F and H). Including the previously estimated affinity constants *K*_3_ and *K*_4_ in the set of fitted model parameters increased model performance (Supplementary Fig. S7 and Supplementary file Supp_table_1), but still failed to account for the observed EC_50_ variations with respect to *γ*_*c*_. This suggested that the structure of model I needed to be modified if it was to reproduce these experimental EC_50_ variations. To this end, we explored how an hypothetical allosteric change induced by JAK3 binding to *γ*_*c*_ could limit binding of IL-7R*α* to *γ*_*c*_ and the formation of the signaling IL-7 receptor complex (see Supplementary section C.2 and Fig. S8). We found that this could result in an increase of the IL-7 EC_50_ for over-abundant *γ*_*c*_. As a result, model performance, reproducing the experimental EC_50_, increased (Supplementary file Supp_table_1 and Supplementary Fig. S8), but the observed variation of EC_50_ for low and intermediate *γ*_*c*_ abundances could not be reproduced for any set of parameters. Hence, allostery cannot account for the measured change of EC_50_ with respect to *γ*_*c*_ expression.

To further explore the imprint of *γ*_*c*_ abundance on cytokine responses, we repeated our experimental measurements and analysis for the STAT5 phosphorylation responses triggered by IL-2 and IL-15 stimulation. In both cases, the effect of *γ*_*c*_ on the amplitude of response to the respective cytokine was nearly identical to that seen following IL-7 treatment (Fig. S4 in Supplementary). In light of this experimental result, the trimeric receptor model (SRLK with *n* = 3) proposed in Ref. (13), with similar amplitude to that of model I, seems to be a good candidate to characterize the IL-2/IL-2R and IL-15/IL-15R systems. Most interestingly, *γ*_*c*_ abundance did not significantly affect EC_50_ values for either IL-2 or IL-15. Thus, while the behavior of the amplitude seems to be similar across several cytokine-receptor systems, the sensitivity of the cellular response, as measured by EC_50_, requires further cytokine-specific experimental and theoretical analysis.

### Specific competition for the formation of the complete IL-7 receptor can account for the EC_50_ increase in the IL-7 response (model II)

Our previous hypothesis of allostery, the existence of differences in binding caused by interactions between the components of the IL-7 receptor complex, did not lead to a mathematical model which could reproduce the experimental data. Thus, we considered the addition of external components to the model (model II). We reasoned that, since the *γ*_*c*_ chain is a component of other cytokine receptors (8), these receptors could limit the amount of *γ*_*c*_ available to form IL-7 receptor complexes, and in turn, decrease IL-7 signaling. To explore this hypothesis we included a putative “decoy” subunit chain (called *x*), that could bind the *γ*_*c*_ chain (and the *γ*_*c*_:JAK3 complex), and sequester it away from IL-7R*α* (see Fig. 3A-C); for instance, *x* could be other surface receptors (such as IL-2R*β* or IL-4R*α*). Further work will be required to identify a suitable candidate for *x* (see section E for an extended discussion). Model II behaves similarly to model I when considering the IL-7 response amplitude (Fig. 3D-E), as expected from the algebraic analysis conducted in Ref. (13), which shows that the amplitude of both models is the same. This guarantees that model II reproduces the experimental amplitude as accurately as model I. As expected, the number of signaling and “dummy” complexes, as well as the number of decoy complexes (with or without JAK3) are limited by their less abundant component (see Fig. 2E). Also, receptor complexes containing the decoy subunity chain *x* are formed only when the number of *γ*_*c*_ exceeds the number of IL-7R*α* chains (*i*.*e*., when the number of *γ*_*c*_ is no longer limiting).

**Fig. 3.**
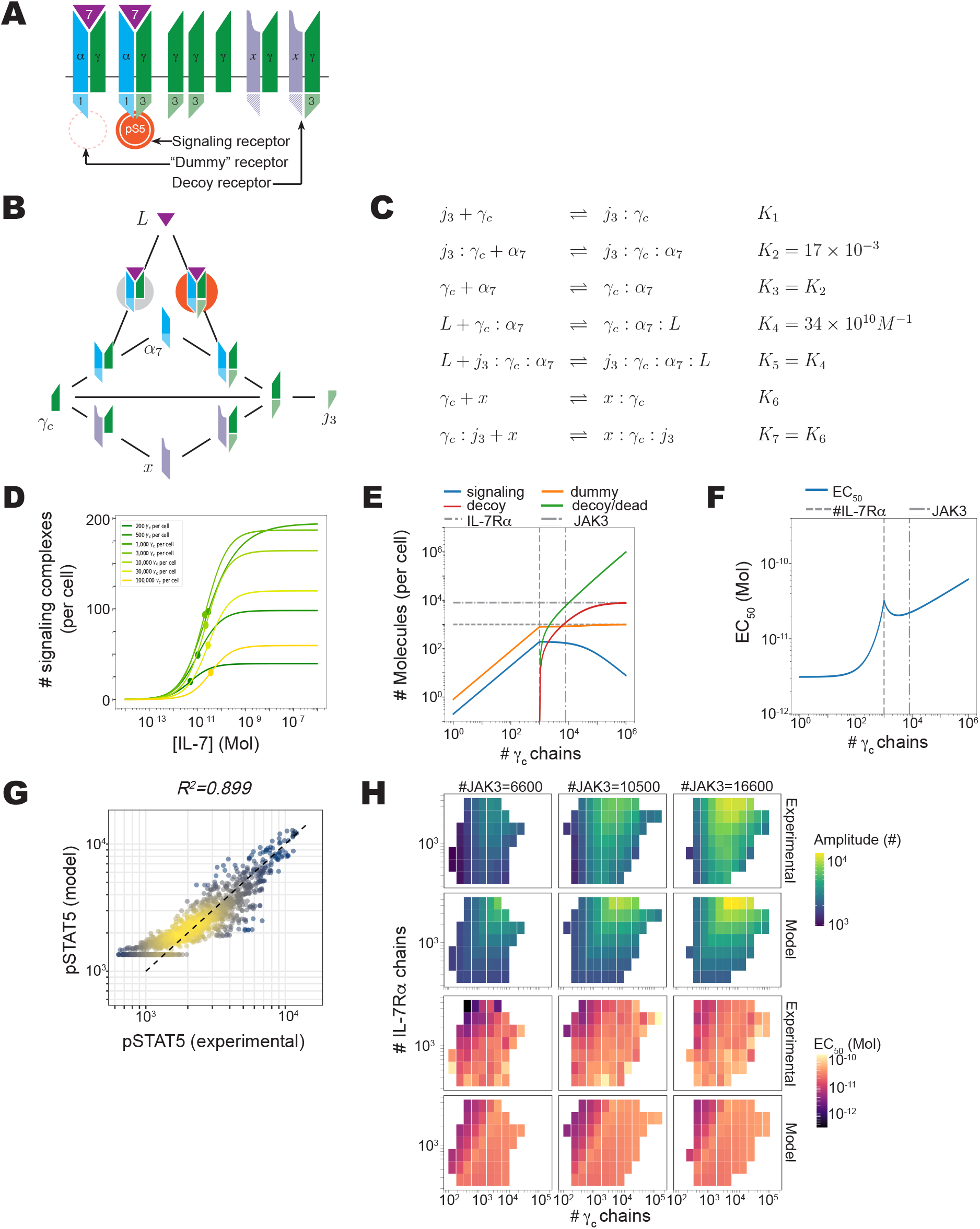
Model II augments model I with a decoy receptor to account for an increased EC_50_, as well as the loss of pSTAT5 response at high levels of *γ*_*c*_. A) Cartoon of the formation of IL-7/IL-7 receptor complex. This model adds a decoy receptor that sequesters *γ*_*c*_. B) Sketch of the biochemical reactions associated with the formation of the IL-7/IL-7R complex. C) Chemical equilibrium for the reactions depicted in B (*j*_3_: JAK3, *α*_7_: IL-7R*α, L*: IL-7, *x*: decoy receptor). *K*_*i*_ are affinity constants for *i* = 1, …, 7. D) Modeled pSTAT5 in response to increasing doses of IL-7 (for different abundances of *γ*_*c*_). E) Amplitude, *i*.*e*., number of signaling (blue) or “dummy” complexes (orange), for increasing number of *γ*_*c*_ chains and for excess cytokine. F) EC_50_ for pSTAT5 in response to IL-7 for increasing levels of *γ*_*c*_. Model II agrees with the experimental results when compared to model I. G) Model II improves predictions (compared to model I) for pSTAT levels across all IL-7 concentrations, as well as across all cellular subpopulations. H) Model II accounts well for the (IL-7R*α, γ*_*c*_) dependence of the pSTAT5 amplitude and EC_50_ (theoretical map to be compared to experimental results). Theoretical results correspond to simulations of model II with best-fit parameters corresponding to the entry “model_II” in Supplementary file Supp_table_1.

The EC_50_ of model II, as is the case in model I, does not depend on JAK3 abundance (as demonstrated in (13)). However it depends on the abundances of *γ*_*c*_ and *x*. More specifically, for a fixed abundance of IL-7R*α*, the EC_50_ increases with the abundance of the *x* chain, and this effect vanishes as *γ*_*c*_ abundance increases (see Supplementary Fig. S5). Hence the presence of the *x* subunit chain provides a way to modulate the EC_50_. Still, a constant amount of *x*, or an amount proportional to the *γ*_*c*_ abundance is not sufficient to reproduce the experimental EC_50_ dynamics. However, when we conjecture that the abundance of the unknown species *x* (*N*_*x*_) increases with that of *γ*_*c*_ (*N*_*γ*_) according to a power-law relation, such that 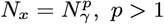, the EC_50_ of model II increases with *γ*_*c*_ (see Supplementary Fig. S6). Moreover, this increase is observed even for low and intermediate *γ*_*c*_ levels, in agreement with our experimental observations.

In this last setting, model II could reproduce both the experimental EC_50_ and the maximum pSTAT5 across subpopulations with different IL-7R*α*, JAK3 and *γ*_*c*_ abundances (Fig. 3H). As a result, a better correlation between theoretical and experimental pSTAT5 levels across all cytokine concentrations was achieved (Fig. 3G). Importantly, the performance of model II was only slightly reduced when the parameters *K*_3_ and *K*_4_ were held fixed at, or constrained to be close to their previously estimated values (see Supp_table_1 and Supplementary Fig. S9), suggesting a consistency between this model and our previous work.

### General insight on the variability and robustness of receptor-ligand signaling systems

Our study demonstrates that the increased expression of a given IL-7 receptor component may paradoxically reduce the amplitude and increase the EC_50_ of the response to IL-7. In this section, we comprehensively explore the impact of varied abundances of receptor onto signal transduction. We generalize models I and II for five different configurations: hetero- or homo-dimeric receptors, each either requiring a downstream kinase or possessing intrinsic kinase activity, and monomeric receptors (Fig. 4A). We note that the hetero-dimeric receptor model with downstream kinase is the configuration of model I. For every receptor system we considered the up/downregulation of a “primary” receptor chain, equivalent to the *γ*_*c*_ subunit chain in our experiments. For the homo-dimeric or monomeric receptors, this is the only receptor chain. For a hetero-dimeric receptor, we set the median abundance of the secondary receptor chain to 10^3^ proteins per cell, by analogy with IL-7R*α* in our system. If a downstream kinase is required, we assumed there is only one kinase (*e*.*g*., JAK3 in the IL-7 receptor system), and set its level to 10^4^ proteins per cell. Finally, we took the (no allostery) affinity constant values described in Fig. 2 (see Supplementary Table S3 and section F): *K*_1_, binding of downstream kinase to primary receptor chain, *K*_2_, binding of primary to secondary (or primary to primary, in the case of homo-dimeric receptors) receptor chain, with or without downstream kinase, and *K*_3_, binding of ligand to the receptor complex, with or without downstream kinase. We used the analytic method presented in Ref. (13) to derive the amplitude and EC_50_ for each of these models when possible. The results are summarized in Supplementary Table S4. We also analysed the global sensitivity of the amplitude and EC_50_ using numerical and machine-learning methods (see Supplementary section G). Our theoretical results and numerical exploration indicate a striking difference for the effect of primary receptor abundance depending on the receptor arrangement:

1. When the primary chain is the only one in the receptor (monomeric or homo-dimeric RTK), increasing the abundance of that chain proportionally increases the amplitude, as expected from Supplementary Table S4. Thus, a 10^2^-fold increase in expression of the primary chain leads to a 10^2^-fold rise in amplitude for the monomeric and a 50-fold rise for the homo-dimeric RTK. This results in a large potential for enhanced signaling (Fig. 4B).
2. The less abundant chain always determines the theoretical maximum of the amplitude. However, this maximum is not attained for receptors with downstream kinase due to the formation of “dummy” complexes (Fig. 4B).
3. Receptors with downstream kinase present a loss of signaling when the number of primary chains increases, thus, rendering the tuning of the response by the primary chain abundance more complex than RTKs (Fig. 4B).
4. The affinity constants of the biochemical reactions only play a minor role in the value of the amplitude. However, they have greater relevance in the EC_50_ (Fig. S10 in Supplementary).

**Fig. 4.**
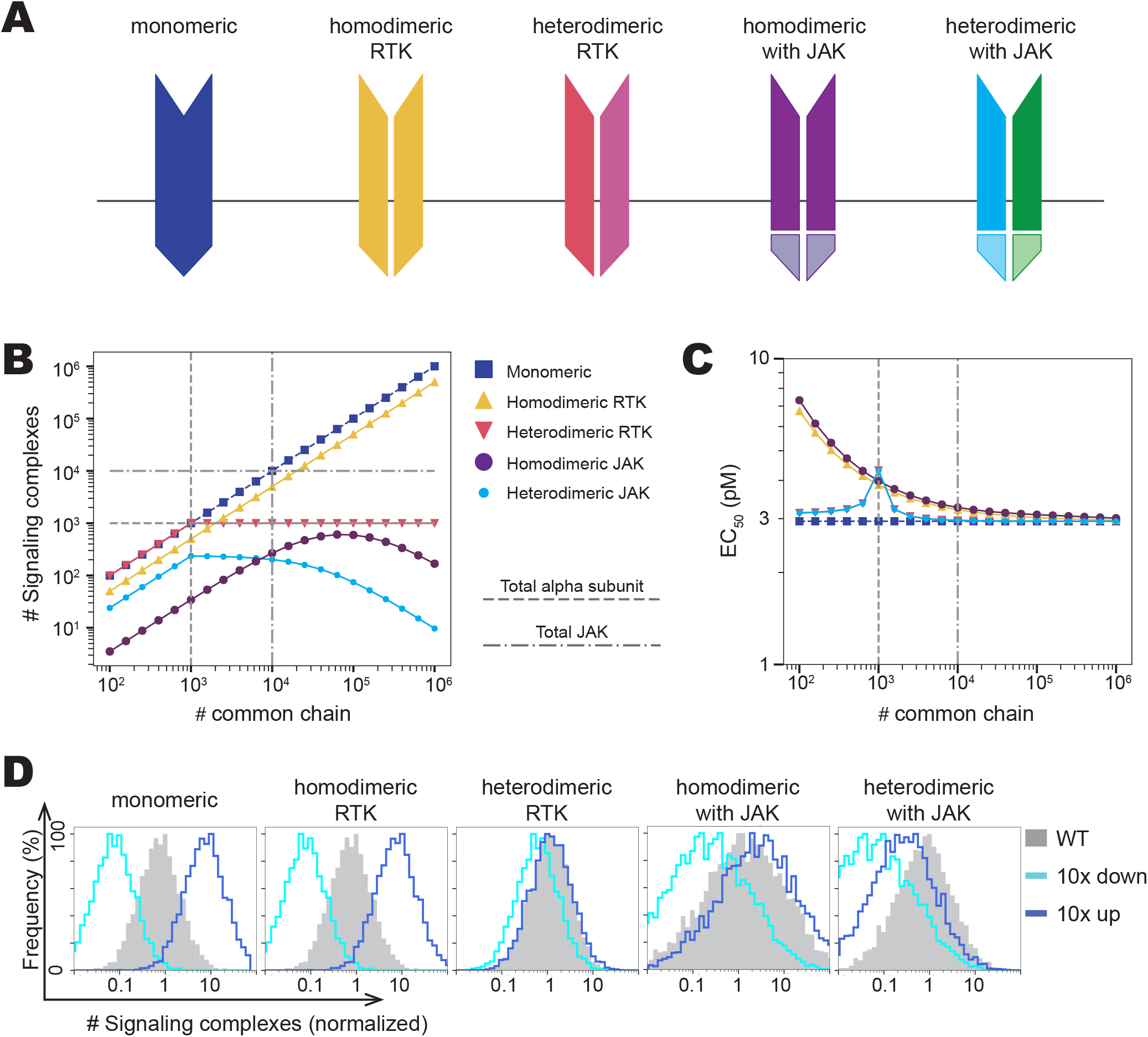
Different architecture of ligand-receptor systems leads to a range of signaling behaviour. A) Receptors might come in five configurations (monomeric, and homo-dimeric or hetero-dimeric, with/without endogeneous kinase). B) Each configuration results in a different relation between the maximal number of signaling receptors (signaling complexes) and the copy number of the primary chain when ligand is in excess, as well as C) EC_50_. D) Distribution of signaling complexes for each receptor configuration when one receptor is up/downregulated by 10-fold.

We then explored the functional implications of the five receptor configurations aforementioned, in terms of signal robustness and variability. Indeed, our experimental studies highlighted the large variability in the abundance of signaling components (receptors, kinases, phosphatases, etc.), even within isogenic population of cells (4, 5). In fact, this variability drives some phenotypic heterogeneity (4, 5, 15, 16). We simulated cellular populations for each of the arrangements, with the distribution of each protein described by a lognormal distribution with a CV of 0.25 (Fig. 4D). The abundance of the secondary receptor chain (if present) was set to have a median of 10^3^ and downstream kinase a median of 10^3.5^, respectively. We then simulated populations with a 10-fold up/downregulation of the primary chain, with a median of 10^4^ for the wild type.

Homo-dimeric receptors with associated kinases (RTKs) or monomeric receptors had the greatest variability of amplitude, matching the distribution of the receptor exactly. Homo-dimeric receptors requiring downstream kinases and hetero-dimeric receptors with associated kinases (RTKs) had the least variability, maintaining amplitude at a moderately high level. Hetero-dimeric receptors requiring downstream kinases had larger variability, but had lower EC_50_ when upregulating or downregulating the primary chain. These results indicated that the composition of a receptor’s signaling core could have strong effects on the variability of cellular responses to extra-cellular ligands. The *γ*_*c*_ family’s arrangement tends to allow for large variability, but leads to lower amplitudes. Hetero-dimeric receptors with intrinsic kinase ability, such as the receptor for insulin, IGF-1, or homo-dimeric receptors with downstream kinases, such as the IL-6 receptor, tend to have intermediate but consistent amplitude. Homo-dimeric receptors with associated kinase activity, such as EGFR, are highly variable in EC_50_ and their amplitude does not saturate (17, 18) (Figs. 4B and C).

We also computed the EC_50_ for each of the receptor configurations (Fig. 4C). We observed that the EC_50_ of the hetero-dimeric models are the same (see Table S4 in Supplementary), and the EC_50_ of the homo-dimeric models are similar (Fig. 4C). This led us to conjecture that the variability of the EC_50_ with the abundance of the primary chain depends on the manner the extra-cellular receptor is built and is independent of its intra-cellular activity. Further experimental work will be required to identify a molecular mechanism to test such conjecture. We note that the EC_50_ of the model in Fig 3 which, this time, captures the experimental behavior, is also independent of the kinase activity of the receptor. We also observe that the EC_50_ for all receptor configurations tends to a constant for large values of *γ*_*c*_ abundance. This value is the dissociation constant of the ligand to the receptor 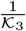 (see Supplementary Table S4).

## Conclusion

We conjecture that the requirement for abundance balance between kinases and receptor subunit chains provides built-in protection against aberrant activation, as observed during oncogenic transformation (17). Robust input/output response does limit functional variability but also protects against enhancement of receptor signaling and its dysregulated accumulation of downstream survival signals, such as Bcl-2. Of course, many additional molecules (or proteins) and biochemical reactions are required to proceed from upregulation of a receptor chain to upregulation of cellular survival signals, not least of which is ligand availability. Nonetheless, the examination of gene expression data from 4,087 tumors sequenced as part of The Cancer Genome Atlas (TCGA) partially validated our conjecture. EGFR was upregulated in 28% of analyzed tumors (compared to healthy counterparts), whereas *γ*_*c*_ was upregulated in only 11%. Furthermore, of the 11% of cases in which *γ*_*c*_ was upregulated, 78% of those also included upregulation of a secondary *γ*_*c*_ family signaling chain, such as IL-4R*α* or IL-2R*β*. Interestingly, IL-7R*α* was upregulated for 24% of the analyzed tumors compared to normal cells. This could reflect different factors, such as the fact that IL-7R*α* was the chain of limiting abundance in our system and/or that IL-7 is ubiquitously available at low levels, from secretion by endothelial cells (19). While increased immune infiltrates could potentially be responsible for the upregulation of lymphocyte-associated receptors in the TCGA tumor samples, the frequent expression in non-hæmatopoietic cancer cell lines (20), and confirmed *in situ* expression of IL-2R*α*, IL-2R*β* and *γ*_*c*_ proteins in breast tumor epithelial cells (21), suggests that expression of cytokine receptors may play a role in non-lymphoid tumorigenesis. We also note that simply quantifying upregulation does not account for the possibility of activating mutations, such as the I241–V253 reported in the IL-7R*α* trans-membrane domain, which allows for ligand-independent activation (22). Hence, a broader analysis tracking receptor upregulation in T or B cell lymphomas over time will provide much insight and validation of the impact of signaling core composition on oncogenicity. Our study, demonstrating that the quantitative effects of receptor chain upregulation can be vastly different, depending on the elements of a receptor’s signaling core, provides a theoretical and quantitative framework with which to interpret the potential functional significance of receptor up/downregulation during lymphocyte differentiation (23, 24), oncogenesis (1) or drug treatment (25).

## Materials and Methods

See Supplementary Information.

## ACKNOWLEDGMENTS

This project has received funding from the European Union’s Horizon 2020 research and innovation programme under the Marie Skłodowska-Curie grant agreement number 764698 (LS and CMP). This work was partially supported by the intramural research program of the NCI (GAB).

