## Supplementary Methods and Figures for "Tuning of cytokine signaling through imbalanced abundances of receptors and kinases"

### Supporting Information Appendix (SI).

#### 1. Materials and experimental methods

**A. Reagents and antibodies.** Recombinant mouse IL-7 and IL-2 were from eBioscience (San Diego, CA). For flow cytometry analysis, monoclonal antibodies against CD4 (clone RM4-5 conjugated with Alexa-700), against IL-2R$\alpha$ (clone PC61.5 conjugated with PE-Cy7 or with PE), against IL-7Ra (clone eBioSB/199 conjugated with PE) were from eBioscience (San Diego, CA). Antibody against JAK3 (clone ) was from Santa Cruz BioTechnology (SCBT, Santa Cruz, CA). Antibody against pY694-STAT5 (clone C11C5) was from Cell Signaling Technology (Danvers, MA).

**B. Media.** All experiments were performed in complemented RPMI medium (prepared by the Media Facility at MSKCC), which consists of RPMI 1640 supplemented with 10% heat-inactivated fetal bovine serum, 2 mM L-glutamine, 10 mM HEPES (pH 7.4), $10^{-1}$ mM non-essential amino acids, 1 mM sodium pyruvate, $10^2$ $\mu$g/ml of penicillin, $10^2$ $\mu$g/ml of streptomycin, and 50 $\mu$M $\beta$-mercaptoethanol. All cell cultures were maintained in an incubator at 37°C with 5% CO2.

**C. Cell culture.** 5C.C7 TCR transgenic, Rag2$^{-/-}$IL-2$^{-/-}$ CD4 T cells were isolated and expanded as described previously (1). Briefly, cells were expanded with irradiated B10A CD3$^{-/-}$ splenocytes pulsed with 1 $\mu$M MCC K5 peptide: ANERADLI-AYFKAATKF (GenScript, Piscataway, NJ). IL-2 (10 nM) was given at one and four days post-stimulation, then every three days for maintenance of cultures. Cells were used between day 5 and 20 of culture, unless otherwise noted. Live cells were purified prior to analysis by separation on a Ficoll gradient (GE, Piscataway, NJ).

**D. Cell transfection to overexpress $\gamma_c$.** Primary T cell blasts were transfected with plasmids expressing $\gamma_c - IRES - GFP$ or $GFP$ alone (as a negative control), using a Nucleofector protocol (Lonza, Walkersville MD). Cells were prepared by mixing 5 million cells with 10 ng of plasmid in Nucleofector solution, incubated for 20 min at room temperature, transfected in the Nucleofector per manufacturer's protocol, incubated again for 30 min at room temperature, added to 5 mL of Nucleofector medium augmented with recombinant human IL-2 (final concentration 1 nM) and placed in the incubator. Cells were typically used 24 hr after transfection.

**E. Cytokine titration.** Before measuring cytokine responses, bound IL-2 was stripped from cell surface by a 2 minute incubation in $10^{-1}$ M glycine buffer equilibrated at pH 4.0. After two washes in RPMI, cells were rested for 30 minutes at 37°C. Cytokine titrations were prepared in 96-well V-bottom plates on ice. Cells ($10^5$ per well) were incubated with cytokines for ten minutes at 37°C before processing for flow cytometry.

**F. Flow cytometry.** Cells were prepared for single cell staining as follow. After stimulation, cells were first fixed for 10 minutes on ice in 1.6% paraformaldehyde followed by permeabilization on ice in 90% methanol. Cells were then washed twice in fluorescence-activated cell sorting (FACS) buffer (PBS with 4% FCS and 0.1% sodium azide) and stained intra-cellularly for phospho-STAT5, followed by incubation with labeled secondary antibody and conjugated antibodies to receptors (CD4 and IL-2R$\alpha$). All staining was performed at room temperature in FACS buffer. Before flow cytometric acquisition, cells were stained with 5 $\mu$M 4',6-diamidino-2-phenylindole (DAPI; Sigma-Aldrich, St. Louis, MO) for exclusion of dividing (G2) and apoptotic (sub-G1) cells. Flow cytometry acquisition was performed on a LSR-II cytometer (BD Biosciences, San Jose, CA).

**G. Receptor abundance calibration.** A standard curve correlating fluorescence, as measured by flow cytometry, to PE molecules was calculated using Bangs Laboratories (Fishers, IN) Quantum R-PE MESF beads. "Corrected median fluorescence intensity" (cMedFI) values were calculated by subtracting the median fluorescence of unstained beads from the median fluorescence value of each PE standard. A linear regression of log cMedFI to log PE bead MESF values was used to establish a standard curve. To calibrate typical abundances of receptor chains, cells were stained with saturating concentrations (as established by titration) of the respective PE-labeled antibodies, or a nonspecific PE-labeled rat IgG control. Corrected median fluorescence was calculated by subtracting the median fluorescence of the isotype control from the median fluorescence of each antibody staining. The calibration curve was used directly to calculate number of molecules from the corrected median fluorescence (Fig. S1), using a ratio of one PE molecule per antibody, as specified by the manufacturer (eBioscience or BD Biosciences).

**H. JAK3 quantification.** JAK3 abundance was first quantified in cells stained only with anti-JAK3 antibody. Fixed and permeabilized cells were first labeled with anti-JAK3 mouse monoclonal antibody, clone 5H2 (Thermo Scientific, Rockford, IL) at a 1:$10^2$ dilution in FACS buffer, at room temperature for 30 minutes in a 96-well plate. Following primary antibody labeling, cells were washed twice in FACS buffer. In parallel QIFIKIT® beads (Dako, Carpinteria, CA), were aliquoted to a well. The QIFIKIT® kit bead sample contains five populations of beads, coated with defined quantities of mouse monoclonal antibody: 2.1 $\times 10^3$, 1.1 $\times 10^4$, 6 $\times 10^3$, 1.8 $\times 10^5$, and 5.8 $\times 10^5$ per bead in the lot used for these experiments. Both beads and cells were stained with the same 1:$10^2$ dilution of FITC conjugated anti-mouse secondary antibody (Dako) for 30 minutes at room temperature. Following labeling, cells and beads were washed twice with FACS buffer, and analyzed via flow cytometry. Using the mean fluorescence intensity (MFI) values of the beads and the defined number of antibodies, a standard curve was fitted to calculate numbers of mouse monoclonal antibodies from MFI. Applying this calculation to the raw JAK3 fluorescence values measured by flow cytometry, we calculated the absolute number of JAK3 molecules per cell. Since JAK3 was measured in the PE channel of our experiments, we made use of "trans-calibration" (described below) to correlate the measured PE values to the calibrated distribution of JAK3 in the FITC channel.

**I. Trans-calibration.** To translate from protein abundances calibrated as single stains in one channel and measured in multi-parameter experiments in another channel, we have developed a simple method of "trans-calibration" to translate between channels. This method makes the conservative assumption that abundance of protein does not depend on the channel in which it is measured, despite the fact that median fluorescence values vary between channels depending on detector voltage, and even coefficients of variation may change. To translate, this method fits a linear model of the log transformed calibrated distribution's percentiles as a function of the log transformed uncalibrated distribution's percentiles. This model is then applied to the individual cells' measured fluorescence values.

**J. Data processing, binning and fitting of dose-responses.** The complete data set contains 432,380 cells corresponding to eight different experimental conditions (seven IL-7 doses ranging from 10 pM to 10 nM, and the control "no-cytokine" condition), for which the pSTAT5 fluorescence intensity along with the absolute number of JAK3, IL-7R$\alpha$, IL-2R$\alpha$ and $\gamma_c$ molecules were quantified. Calibrated data were $log_{10}$-transformed and subdivided into bins corresponding to different total numbers of JAK3, IL-7R$\alpha$ and $\gamma_c$ proteins per cell, denoted as #JAK3, #IL-7R$\alpha$ and #$\gamma_c$, respectively. For the #IL-7R$\alpha$ and #JAK3 variables, bins of width 0.2 were created in the interval ranging from the 5% to the 95% quantile of $log_{10}$-transformed values. For the #$\gamma_c$ variable, bins of width 0.25 were created in the entire range (from min to max) of $log_{10}$-transformed values. When the total number of IL-2R$\alpha$ proteins per cell (denoted as #IL-2R$\alpha$) was considered as an additional binning variable (Fig. S2), bins of width 0.25 were created in the interval ranging from the 5% to the 95% quantile of $log_{10}$-transformed values. In all binning scenarios, only bins containing at least five cells for each experimental condition were selected for further analysis. For each bin, the pSTAT5 (denoted $p_5$) response to IL-7 stimulation was fitted to a Michaelis-Menten function with a background term, as follows

$$p_{5_{\text{exp}}} = \beta_1 + \beta_2 \frac{L}{(EC_{50} + L)}, \quad [1]$$

where $L$ denotes the extra-cellular IL-7 concentration. Values for $\beta_1$, $\beta_2$, and $\mathrm{EC}_{50}$ were estimated using a non-linear least square analysis. We minimized the sum of differences between predicted and measured mean $log_{10}$-transformed pSTAT5 values, weighted by the standard error of the mean. From this analysis, we selected 172 bins for which convergence towards a minimum was achieved, and for which the estimated $\mathrm{EC}_{50}$ parameter was higher than $10^{-13}M$ with a 95% confidence interval within $[10^{-20}M, 10^{-8}M]$.

#### 2. Mathematical modeling

**A. Variability of the affinity constant $K_D$ for cytokine binding, as revealed by varied relations between $EC_{50}$ and amplitude for different $\gamma_c$ cytokines..** We present here a coarse-grained biochemical model of STAT phosphorylation upon cytokine engagement with its receptor on the surface of cells, in order to derive the dose-response relation of phosphorylated STAT ($pSTAT$), as a function of the cytokine concentration. Our goal is to highlight how the maps of $EC_{50}$ and amplitude for different receptor abundances challenge a simple biochemical model of cytokine signaling.

A cartoon of our model is presented in Figure S3A-B. In a nutshell, cytokines engaged their receptor on the surface of cells. Ligand-bound receptors act as kinases that phosphorylate STAT molecules. We assume a constant dephosphorylation rate.

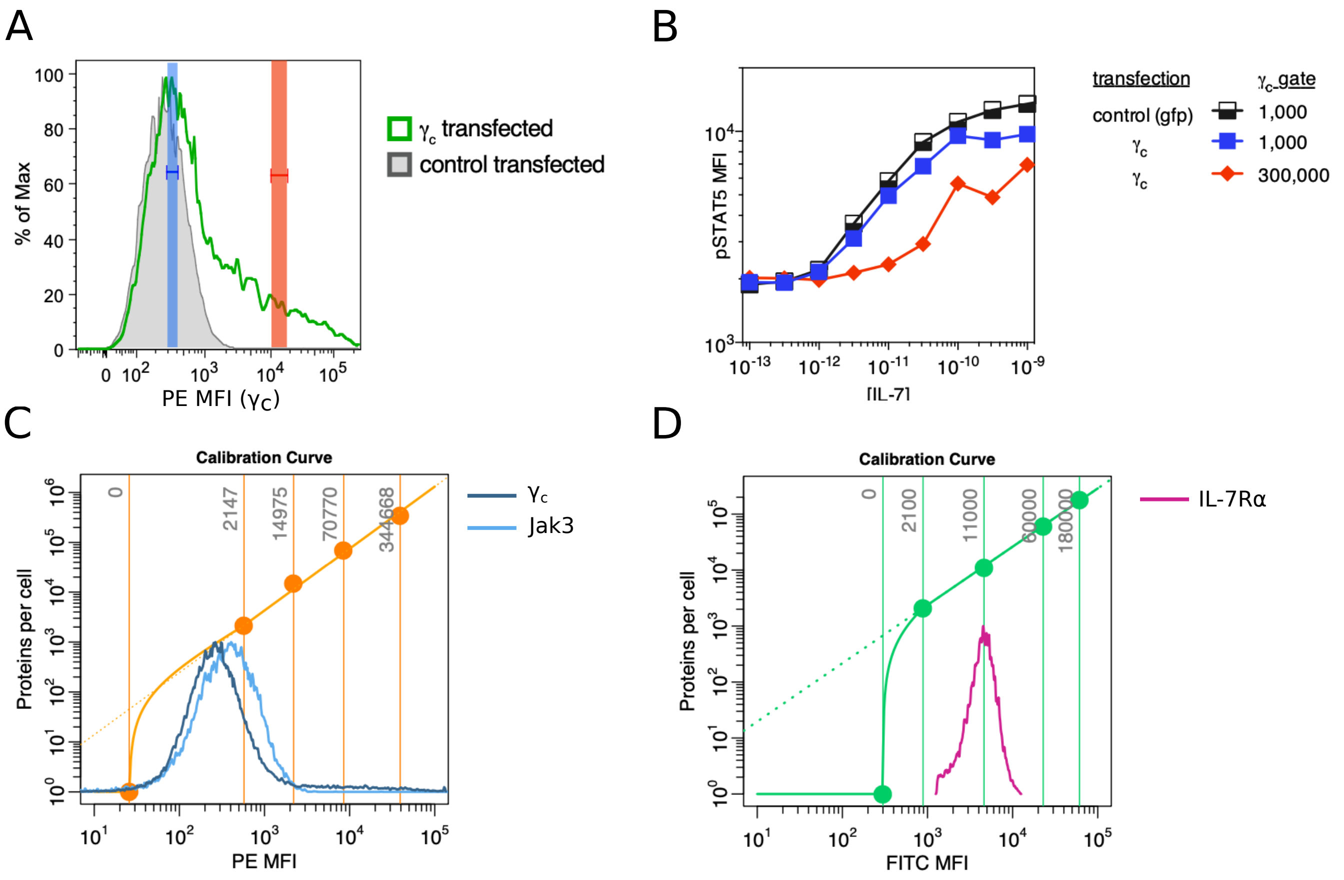


**Fig. S1.** Characterization of the altered response to IL-7 for high abundance of $\gamma_c$. A) Distribution of the expression levels of $\gamma_c$ on the surface of T cell blasts, after transfection of a plasmid expressing $\gamma_c$, measured by antibody staining and flow cytometry. The two colored vertical bands (blue and red) define two subpopulations expressing low and high levels of $\gamma_c$, respectively, within the transfected cells. B) Dose-response of STAT5 phosphorylation for increasing doses of IL-7 for different levels of $\gamma_c$: (black) control cells expressing GFP; (blue and red) cells transfected with $\gamma_c$ expressing plasmids, and gated for low and high levels, respectively. C) Quantitation of the average abundance of JAK3 and $\gamma_c$ in primary cells; the orange point and line are calibration beads; the histograms are the distribution of molecules measured by antibody staining and flow cytometry. D) Quantitation of the average abundance of IL-7R$\alpha$ (performed as in C).

Based on our previous experimental observations (1), this signal transduction cascade can be modeled at steady-state. Many molecular details of this signaling pathway (*e.g.,* conformational changes of activating receptor-associated kinases JAK) are being neglected for the sake of simplicity, without compromising the generality of our results.

If we follow classical biochemical kinetics, we can describe the time evolution of pSTAT as follows:

$$\frac{d}{dt}[pSTAT] = k_{cat}[STAT \cdot R^*] - k_{dephos}[pSTAT] \; , \qquad [2]$$

where we have made used of the Michaelis-Menten model of enzymatic activity to approximate

$$[STAT \cdot R^*] = \frac{k_+}{k_- + k_{cat}}[STAT][R^*] \; . \qquad [3]$$

In the previous equation we have abused notation to denote by $[R^*]$ the abundance of engaged/active receptors. We now make use of the fact that the total number of STAT molecules is conserved ($S_0 = [STAT] + [pSTAT]$) and we assume steady-state

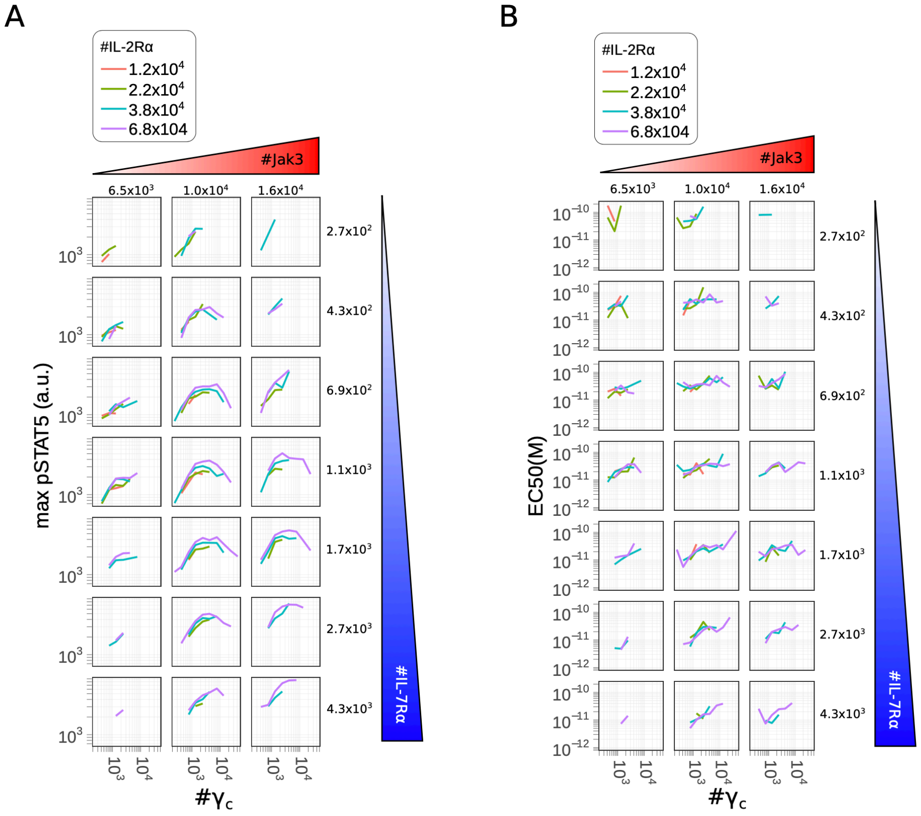


**Fig. S2.** A, B – Representation of the maximum pSTAT5 intensity (A) and IL-7 $EC_{50}$ (B) as a function of the number of $\gamma_c$ molecules per cell. Each point corresponds to a bin, a subpopulation of cells whose number of $\gamma_c$, IL-7R$\alpha$, JAK3 and IL-2Ra molecules per cell lie within a given distance from the bin center (see methods), as indicated on top and left for JAK3 and IL-7R$\alpha$, respectively, and in the figure key for IL-2R$\alpha$. For each bin, the maximum pSTAT5 and $EC_{50}$ are estimated from a fit of the experimental pSTAT5 values with a non-cooperative Hill function (see methods).

conditions in Eq. (2) to write

$$[pSTAT] = \frac{k_{cat}}{k_{dephos}}[STAT \cdot R^*] = \frac{S_0}{1 + \frac{k_{dephos}}{k_{phos}[R^*]}} \ , \qquad [4]$$

with $k_{phos} = \frac{k_+ k_{cat}}{k_- + k_{cat}}$. In order to obtain the number of engaged/active receptors, $[R^*]$, we use the chemical equilibrium equation combined with the conservation of total receptor numbers. We have

$$[R^*] = \frac{1}{K_D}[cytokine][R] \ , \quad \text{and} \quad R_0 = [R] + [R^*] \ , \qquad [5]$$

where $R_0$ is the total number of receptors. We then obtain

$$[R^*] = R_0 \left(1 + \frac{K_D}{[cytokine]}\right)^{-1} \ , \qquad [6]$$

and the following equation for the pSTAT dose-response:

$$[pSTAT] = \frac{S_0}{1 + \frac{k_{dephos}}{k_{phos} R_0}\left(1 + \frac{K_D}{[cytokine]}\right)} \ . \qquad [7]$$

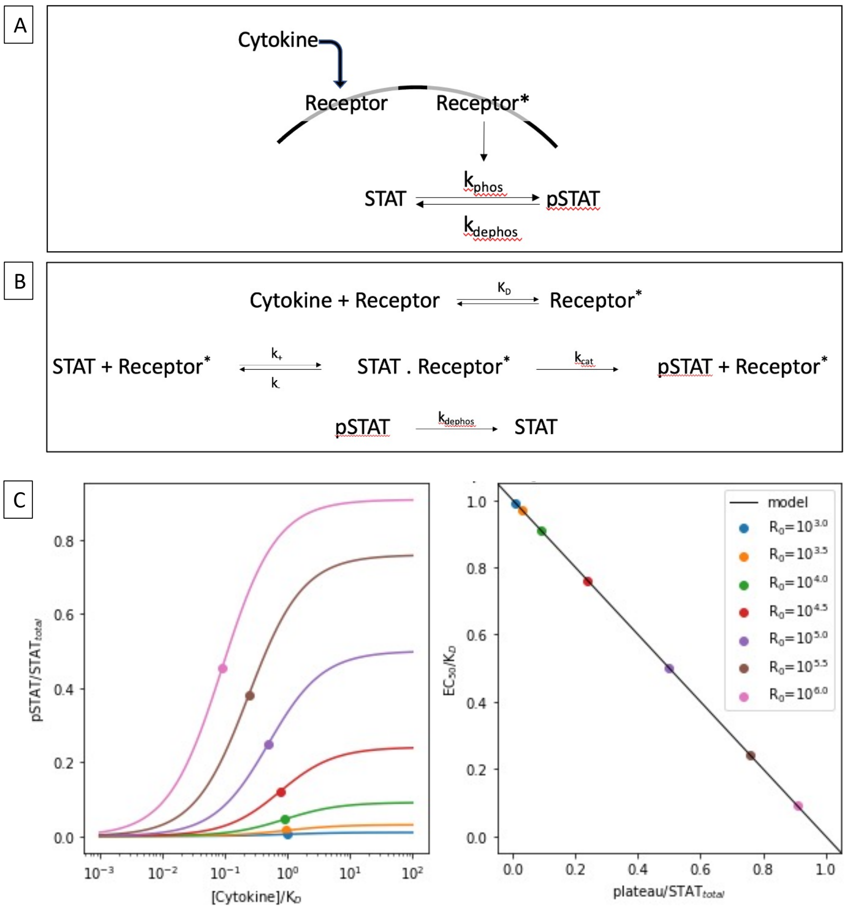


**Fig. S3.** Simple biochemical model to relate $P_\infty$ and $EC_{50}$ for the cytokine dose-response curve. A) Sketch of the biochemical reactions associated with cytokine binding and STAT activation. B) Corresponding biochemical reactions and associated rates. C) Computed dose-response curves for pSTAT as a function of the cytokine concentration (plotted for different receptor abundances $R_0$ (left), and predicted linear relation between $\mathrm{EC}_{50}$ and $P_\infty$ (right). Such linear relation is not consistent with our experimental results (when comparing STAT5 phosphorylation for IL-2, IL-7 and IL-15, hence a simple model of cytokine binding to its receptor must be ruled out (see Figure S4).

These dose-response curves are plotted in Figure S3C (left) for increasing levels of receptor abundances, $R_0$. From the previous equation we can derive the amplitude, $P_\infty$, of the dose-response curve:

$$P_\infty = pSTAT\left([cytokine] \to +\infty\right) = \frac{S_0}{1 + \frac{k_{dephos}}{k_{phos} R_0}} \,, \quad [8]$$

and the $EC_{50}$, defined as the cytokine concentration which satisfies $pSTAT(EC_{50}) = \frac{1}{2}P_\infty$. This value is given by

$$EC_{50} = K_D \left(1 + \frac{k_{phos} R_0}{k_{dephos}}\right)^{-1} \,, \quad [9]$$

which leads to a linear relation between $EC_{50}$ and the amplitude

$$EC_{50} = K_D \left(1 - \frac{P_\infty}{S_0}\right) \,. \quad [10]$$

In particular, we obtain two extreme regimes of behaviour: for small number of receptors, or when dephosphorylation is much faster than phosphorylation, the amplitude $P_\infty$ tends to zero and $EC_{50}$ tends to $K_D$, the affinity of the cytokine to its receptor; inversely, for large receptor abundances, or when dephosphorylation is much slower than phosphorylation, the amplitude tends to the total number of STAT molecules, $S_0$, and the $EC_{50}$ vanishes.

We presented this simple model to demonstrate that $K_D$, the affinity of a cytokine to its receptor on the surface of cells, cannot be a fixed biochemical constant. When comparing the amplitude ($P_\infty$) of STAT5 phosphorylation, induced by IL-2, IL-7 and IL-15 for T cells, we found similar variability according to their levels of IL-2R$\alpha$ and common $\gamma_c$ receptor chains (Fig. S4). Most likely, the kinase activities of the engaged receptors are larger than the phosphatase activivites, such that $k_{dephos} \ll k_{phos} R_0$, and $P_\infty = S_0$ (at high cytokine concentration, all STAT molecules available get phosphorylated). Consequently, our result, Eq. (10), would predict that $EC_{50}$ for these three cytokines should be proportional to each other (according to their different values of $K_D$). The differences of $EC_{50}$ maps in Fig. S4 are inconsistent with the simple modeling results derived here; hence our initial assumption that $K_D$ is a constant, independent of the different receptor copy numbers in cells, needed to be revisited.

**B. Modeling hypotheses.** In order to make the modeling more tractable, we made experimentally-justified approximations (see Table S1).

First, since STAT5 phosphorylation in response to $\gamma_c$ cytokines reaches steady-state within ten minutes of exposure (2, 3), our model assumes that all biochemical reactions reach a steady-state to calculate the abundance of each complex. This simplification allows us to make use of experimentally-determined equilibrium constants ($K$) to quantify the interaction between receptor components in the membrane (4), rather than kinetic rates for the forward and backward biochemical reactions (on and off rates), which have not been experimentally determined. Second, since the inhibitory effect of $\gamma_c$ was observed for a fixed amount of IL-2R$\alpha$ (Supplementary Fig. S2), we omitted the IL-2 receptor components in our initial model. Third, the parameters defining STAT5 phosphorylation have not been determined. In our previous work, we demonstrated that the

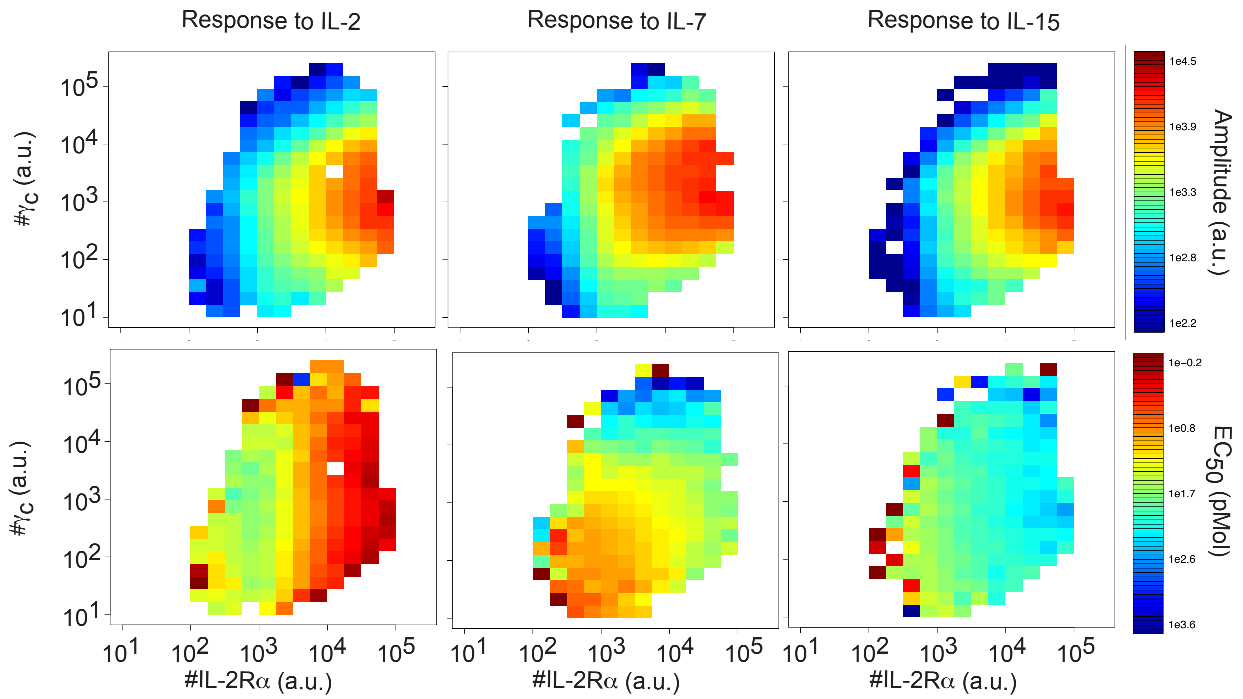


**Fig. S4.** Maps of amplitude $P_\infty$ (top) and EC$_{50}$ (bottom) for the pSTAT5 responses to IL-2, IL-7 and IL-15, for different expression levels of IL-2R$\alpha$ and $\gamma_c$. Note how similar the amplitude values, $P_\infty$, are for different cytokines, while the EC$_{50}$ have very different dependencies with the abundances of IL-2R$\alpha$ and $\gamma_c$. This experimental observation is to be contrasted with a contant $K_D$ model which predicted a simple anti-correlation between $P_\infty$ (top) and EC$_{50}$ (see Supplementary section A and Figure S3)

.

abundance of pSTAT5 correlated linearly with the number of IL-7 bound to the cell surface (1), hence we assumed that the fraction of phosphorylated STAT5 molecules and the number of IL-7R$\alpha$–$\gamma_c$–IL-7 complexes, $\sigma$, follow the affine relation:

$$pSTAT5 = \lambda_1 + \lambda_2 \sigma, \qquad [11]$$

where $\lambda_1$ and $\lambda_2$ are parameters to be determined. Finally, since IL-7R$\alpha$ depends on JAK1 for signaling, a similar balance would exist for these proteins. However, though we tested multiple antibodies, we could not find one suitable for the flow cytometric analysis of JAK1. Consequently, we focused our modeling effort on the effects of IL-7R$\alpha$, JAK3, and $\gamma_c$ abundances, which could be experimentally quantified, and thus, estimated the number of phosphorylated STAT5 molecules as proportional to the number of fully-formed signaling complexes IL-7R$\alpha$–$\gamma_c$–IL-7–JAK3 (see Fig. 2D), calculated from the equilibrium amounts its molecular components.

**C. IL-7R models.** The IL-7R models considered in this manuscript are all sub-models of the following general case, where we considered the biochemical reactions with their affinity constant:

$$\begin{array}{lcll}
j_3 + \gamma_c & \rightleftharpoons & j_3 : \gamma_c & K_1 \\
j_3 : \gamma_c + \alpha_7 & \rightleftharpoons & j_3 : \gamma_c : \alpha_7 & K_2 \\
\gamma_c + \alpha_7 & \rightleftharpoons & \gamma_c : \alpha_7 & K_3 \\
L + \gamma_c : \alpha_7 & \rightleftharpoons & \gamma_c : \alpha_7 : L & K_4 \\
L + j_3 : \gamma_c : \alpha_7 & \rightleftharpoons & j_3 : \gamma_c : \alpha_7 : L & K_5 \\
\gamma_c + x & \rightleftharpoons & x : \gamma_c & K_6 \\
j_3 : \gamma_c + x & \rightleftharpoons & x : j_3 : \gamma_c & K_7
\end{array}$$

where the common gamma chain is denoted by $\gamma_c$, IL-7R$\alpha$ by $\alpha_7$, JAK3 by $j_3$, the unspecified $\gamma$-binding subunit by $x$, and the ligand IL-7 by $L$. A complex formed by two proteins $a$ and $b$ is denoted as $a : b$ or $b : a$. We write $k_i$ for the forward reaction constants (formation of the complex $a : b$ by association of $a$ and $b$), and $q_i$ for the backward reaction constants (dissociation of the complex $a : b$ in $a$ and $b$). Thus, the affinity constants are given by $K_i = \dfrac{k_i}{q_i}$. We will write $[a]$ to denote the concentration of the chemical species $a$ (in number of molecules per cell for $\gamma_c$, $\alpha_7$ and $j_3$, in concentration for $L$). To obtain our mathematical models, we assumed mass action kinetics for the reactions, which means the rates are proportional to the concentration of the reactants. We also assumed that spatial inhomogeneities can be ignored, which allows us to formulate the model in terms of ordinary differential equations (ODEs). We consider the models at steady-state (equilibrium) and assume that the number of $\gamma_c$, $\alpha_7$ and $j$ are conserved. We can then write steady-state and conservation equations from the ODEs,

| Simplification | Benefit | Justification |
|---|---|---|
| Use of equilibrium steady-state approximation, rather than kinetic modeling. | Equilibrium constants ($K_A$) for association of intact trans-membrane receptors have been measured in living cells. | IL-7 signaling reaches equilibrium within 10 minutes of exposure (2). |
| Estimate the abundance of STAT5 phosphorylation as the number of complete IL-7R$\alpha$–$\gamma_c$–IL-7–JAK3 complexes. | Parameters for STAT5 phosphorylation have not been measured. | pSTAT5 correlated linearly with the number of IL-7 bound complexes to the surface of cells (1). |
| Omit JAK1 from the model (rather than requiring JAK1 binding to IL-7 for pSTAT5). | JAK1 abundance could not be experimentally quantified, and would require an additional free parameter. | Low abundance of IL-7R$\alpha$ implies that JAK1 would unlikely be limiting. |
| Only include IL-7 receptor components in model, exclude IL-2 receptor subunits. | Including additional $\gamma_c$ family receptors would add to the complexity of the model. | The impact of varying levels of $\gamma_c$ was detected at low levels of IL-2R$\alpha$, where cross-inhibition would be minimal. |

**Table S1. Summary of model simplifications (or approximations) and their justification.**

which once combined, give a polynomial system that characterizes the model:

$$
\begin{aligned}
0 &= -N_{\gamma_c} + [\gamma_c] + K_3[\gamma_c][\alpha_7] + K_1[\gamma_c][j_3] + K_2K_1[\gamma_c][\alpha_7][j_3] + K_4K_3L[\gamma_c][\alpha_7] + K_5K_2K_1[L][\gamma_c][\alpha_7][j_3] \\
&\quad + K_6[x][\gamma_c] + K_7K_1[x][\gamma_c][j_3], \\
0 &= -N_{\alpha_7} + [\alpha_7] + K_3[\gamma_c][\alpha_7] + K_2K_1[\gamma_c][\alpha_7][j_3] + K_4K_3[L][\gamma_c][\alpha_7] + K_5K_2K_1[L][\gamma_c][\alpha_7][j_3], \\
0 &= -N_{j_3} + [j_3] + K_1[\gamma_c][j_3] + K_2K_1[\gamma_c][\alpha_7][j_3] + K_5K_2K_1[L][\gamma_c][\alpha_7][j_3] + K_7K_1[x][\gamma_c][j_3], \\
0 &= -N_x + [x] + K_6[x][\gamma_c] + K_7K_1[x][\gamma_c][j_3],
\end{aligned} \qquad [12]
$$

where we wrote $N_a$ for the total number of $a$ molecules per cell. The variables are the concentrations at steady-state of the $\gamma_c$, IL-7R$\alpha$, and JAK3 molecules.

We studied the models computationally by solving the resulting polynomial system Eq. (12) with Python or R, for different values of $N_{\gamma_c}$ and ligand concentration, $L$. The number of signaling complexes is given by:

$$\sigma(L, \alpha_7, \gamma_c, j_3) \equiv K_5K_2K_1[L][\alpha_7][\gamma_c][j_3]. \qquad [13]$$

For fitting purposes, the number of signaling complexes is then converted to pSTAT5 fluorescence intensity making use of the equation Eq. (11).

The number of dummy complexes is given by:

$$K_4K_3[L][\gamma_c][\alpha_7]. \qquad [14]$$

The amplitude of the dose-response curve is obtained by considering the function $\sigma$ for high ligand concentration. The $EC_{50}$ of Fig. 2F and Fig. 3F is computed by fitting the dose-response curve (number of signaling complexes as a function of ligand concentration, typically $10^{-15}$ to $10^{-4}$M) to a sigmoid function:

$$f(L) \equiv \frac{\text{amplitude}}{1 + e^{-k(L-EC_{50})}}, \qquad [15]$$

where $k$ and $EC_{50}$ are parameters to be determined. In other cases, the $EC_{50}$ was computed making use of a bisection search.

***C.1. Model I (Fig. 2).*** The first model presented in our study (Fig. 2) does not take into account the $\gamma_c$-binding protein $x$. We also assumed there is no allostery, which means that the affinity constants of the reactions involving the molecule JAK3 are the same as the affinity constants of the same reaction without JAK3. Hence the first model is obtained by setting $K_6 = K_7 = 0$, $K_2 = K_3$ and $K_4 = K_5$. This model, therefore, has a maximum of five free parameters: $K_1$, $K_3$, $K_4$, $\lambda_1$, and $\lambda_2$. The number of free parameters reduces to three when we set $K_3 = 17 \times 10^{-3}$ and $K_4 = 34 \times 10^{10} M^{-1}$, as estimated in our previous study (1).

***C.2. Allostery model.*** We tested the hypothesis that the formation of "dummy" complexes is faster than that of the signaling ones. This results in the monopolization of the $\gamma_c$ chain by non-signaling receptors. We revised our model I to allow for allostery; that is, the affinity constants of the "dummy" and signaling complexes are not equal anymore. The allostery model does not

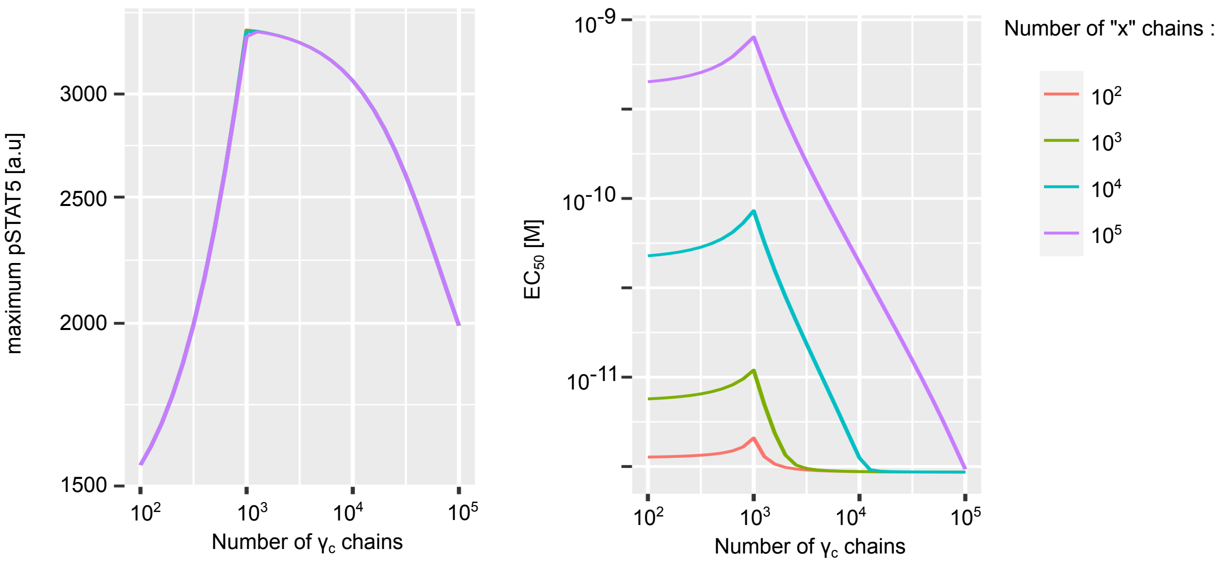


**Fig. S5.** Effect of different abundances of the $\gamma_c$-binding $x$ chain on the maximum pSTAT5 (left) and the $EC_{50}$ (right) for model II. Simulations were performed using $10^3$ IL-7R$\alpha$ chains and $10^4$ JAK3 kinases, and best-fit parameters corresponding to the entry "model_II" in Supplementary file Supp_table_1.

consider the presence of the $\gamma_c$-binding protein $x$, but allows the binding of $L$ (IL-7) to $\alpha_7 : \gamma_c$, and the binding of $\alpha_7$ to $\gamma_c$ to depend on the presence of JAK3. This model, thus, has a maximum of seven free parameters: $K_1$, $K_2$, $K_3$, $K_4$, $K_5$, $\lambda_1$, and $\lambda_2$. The allostery model reproduced the observed amplitude and there exists a set of parameters where the $EC_{50}$ does indeed increase with the abundance of $\gamma_c$ overall. However, the general pattern of the $EC_{50}$ did not agree with the observed experimental evidence: the modeled $EC_{50}$ increased only once the amount of $\gamma_c$ was greater than the amount of IL-7R$\alpha$ (see Fig. S8B), while the experimental $EC_{50}$ seemed to present a regular increase.

***C.3. Model II (Fig. 3).*** Model II, in a similar fashion as model I, does not consider JAK3-induced allostery. In this model however, the cellular abundance of the $\gamma_c$-binding protein $x$ is assumed to scale with the cellular abundance of $\gamma_c$ as follows: $N_x = N_{\gamma_c}^p$. Hence, this model is derived from the general model by imposing $K_2 = K_3$, $K_4 = K_5$ and $K_6 = K_7$. The model has a maximum of seven free parameters: $K_1$, $K_3$, $K_4$, $K_6$, $\lambda_1$, $\lambda_2$ and $p$. Fig. S6 displays the first numerical exploration conducted on the values of $K_6$ and $p$ to assess whether model II could reproduce the observed $EC_{50}$ increase. It shows that $p > 1$ is required to observe an increase in sensitivity.

**D. Fitting of model parameters.** For each model, best-fit parameters were estimated as those minimizing the squared distance, $\Delta^2$, between model and experimental estimates of maximal pSTAT5 and $EC_{50}$ across all 172 bins (obtained as described in Section J). The squared distance $\Delta^2$ was defined as:

$$\Delta^2 = \Delta^2_{EC_{50}} + \Delta^2_{p5_{max}}, \qquad [16]$$

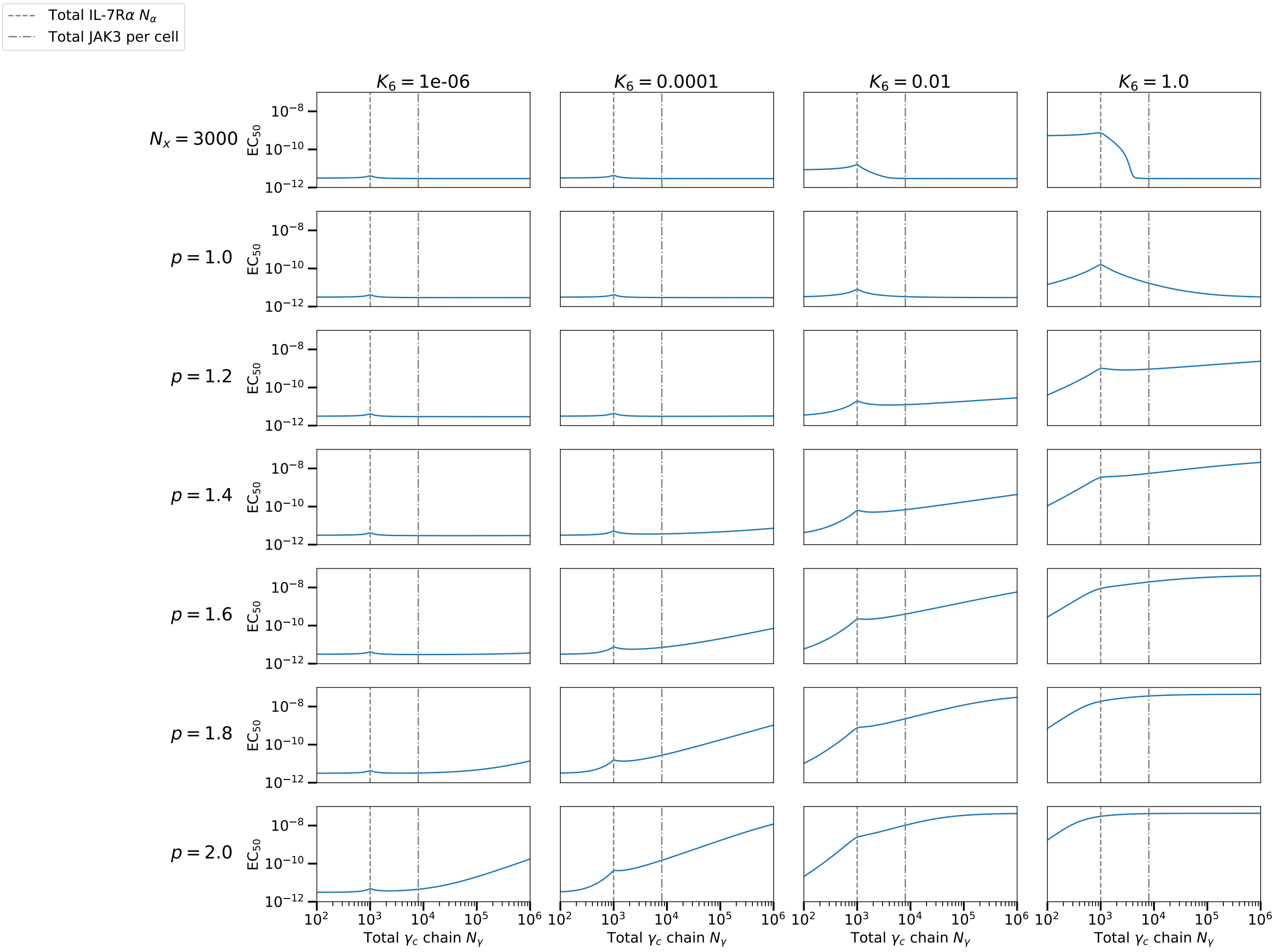


**Fig. S6.** $\mathrm{EC}_{50}$ of model II as a function of the total number of gamma chains, with $10^3$ alpha chains per cell and $8 \times 10^3$ JAK3 molecules per cell, $K_1 = 10^{-4.5}$ and for different values of $p$ and $K_6$. We also explored the case of a fixed ($p = 0$) total abundance of $x$ subunit chain, $N_x$, at $3 \times 10^3$ molecules per cell. We observe an increasing $\mathrm{EC}_{50}$ for $p > 1$. The smaller $K_6$, the larger $p$ has to be to observe an increasing $\mathrm{EC}_{50}$.

with

$$\Delta^2_{EC_{50}} = \sum_{bin} \left[ \log\left(\frac{EC_{50}^{model}[bin]}{EC_{50}^{exp}[bin]}\right) \right]^2, \qquad [17]$$

and

$$\Delta^2_{p5_{max}} = \sum_{bin} \left[ \log\left(\frac{p5_{max}^{model}[bin]}{p5_{max}^{exp}[bin]}\right) \right]^2. \qquad [18]$$

When a prior distribution was considered, model parameters were estimated as those minimizing $\Delta^2 = \Delta^2_{EC_{50}} + \Delta^2_{p5_{max}} + \Delta^2_{par}$, where $\Delta^2_{par}$ quantifies the distance between parameters and their prior estimates:

$$\Delta^2_{par} = \sum_{par} \left[ \log\left(\frac{par}{par_{prior}}\right) \right]^2.$$

The sum runs over the $K_3$, $K_4$, and $\lambda_2$ parameters since prior information was available only for these three parameters. Prior estimates for these parameters were taken from Ref. (1) as $K_{3prior} = 17 \times 10^{-3}$, $K_{4prior} = 34 \times 10^{10} M^{-1}$, and $\lambda_{2prior} = 10$.

Minimization was performed using a Nelder-Mead simplex algorithm. Errors on best-fit parameters were estimated from the covariance matrix defined as the inverse of the Hessian matrix of the minimization function (namely $\Delta^2$) evaluated at the best-fit parameters position.

Best-fit parameters and a summary of the results for the different fits performed can be found in file Supp_table_1. See also Supplementary Fig. S9 for a representation of the different scores assessing fit quality.

**E. Identification of the extra chain $x$ in model II: a discussion.** To quantitatively reproduce the increase of the $EC_{50}$ to IL-7 with increasing $\gamma_c$ abundance, we postulated the existence of a $\gamma_c$-binding protein, $x$, whose abundance would correlate with that of $\gamma_c$. We hypothesized that this $\gamma_c$-binding protein, $x$, could be a subunit chain from a $\gamma_c$ family receptor. Among those candidate subunits, we first considered the beta chain of the IL-2 receptor (IL-2R$\beta$). Using our previous model of IL-2R signaling, the $EC_{50}$ to IL-2 is predicted to decrease with increasing IL-2R$\beta$, and hence with increasing $\gamma_c$. However, experimental evidence indicated that the $EC_{50}$ to IL-2 is largely independent of $\gamma_c$ abundance (Supplementary Fig. S4), hence indicating that the postulated $x$ protein is unlikely to be IL-2R$\beta$. As IL-2R$\beta$ chain is also part of the IL-15 receptor, a similar observation that the $EC_{50}$ to IL-15 did not depend on $\gamma_c$ abundance further refuted the hypothesis that IL-2R$\beta$ could play the role of the $x$ protein.

We next considered the scenario in which the $x$ protein is a subunit of another $\gamma_c$ family receptor, other than IL-2R$\beta$. In this case, our model of IL-2R signaling predicts that the $EC_{50}$ to IL-2 increases with increasing $\gamma_c$ in a similar fashion as observed for the $EC_{50}$ to IL-7. This prediction does not agree with the experimental observation mentioned above that the $EC_{50}$ to IL-2 does not depend on $\gamma_c$ abundance. Hence, the fact that the increase of the $EC_{50}$ with increasing $\gamma_c$ could be observed in response to IL-7 but not to IL-2 or IL-15 stimulation suggests a complex mechanism in which the postulated $x$ protein would act only on the IL-7 receptor.

**F. Models for alternative receptor configurations.** This section analyses the receptor systems described in Fig. 4 which are a combination of a primary chain $\gamma$, a secondary chain $\alpha$, if present, and a JAK molecule $j$, when we include downstream kinase activity. The receptors bind to a ligand, $L$, which is assumed to be in excess. Similarly to the previous models, when a downstream kinase is required, we allow the formation of dummy complexes, the receptor without JAK, and assume no allostery: the affinity constants of the reaction between the ligand and the dummy receptor and the ligand and the signaling receptor are the same, the affinity constant of the reaction between the secondary and the primary chain chain (or the dimerization of the primary chain) is the same whether the $\gamma$ chain is bound to a JAK molecule or not. Thus, we denote by $\mathcal{K}_1$, the affinity constant of the binding between JAK and the primary receptor chain, $\mathcal{K}_2$, the affinity constant of the binding between primary to secondary receptor chain (or the dimerization of the primary chain), with or without downstream kinase, and $\mathcal{K}_3$, the affinity constant of the binding between ligand and the full receptor $\alpha : \gamma$ (or $\alpha : \gamma : j$). Finally, we study the systems at steady-state and assume the conservation of the chains $\gamma$, $\alpha$ and $j$. We write $N_\gamma$, $N_\alpha$ and $N_j$ for the total number of $\gamma$, $\alpha$ and JAK chains per cell, respectively. We write $[X]$ to denote the concentration of the species $X$, and $X : Y$ denotes the complex formed by the species $X$ and $Y$. The polynomial systems describing the models are then derived from the biochemical reaction, as discussed in section C. One needs to pay attention to the factor 2 in the homo-dimeric models. This factor represents the stoichiometric difference between hetero-dimeric and homo-dimeric models. We solved numerically the polynomial systems using Python and plotted the number of signaling complexes at high ligand concentration to obtain the amplitude (Fig. 4B). The dose-response curve (number of signaling complexes as a function of ligand concentration, $L$) can be fitted to a sigmoid equation and the $EC_{50}$ is a parameter of this equation (Fig. 4C).

***F.1. Monomeric receptor.*** The monomeric receptor is composed of a single $\gamma$ chain which binds to the ligand. The biochemical reaction and its affinity constant are

$$L + \gamma \rightleftharpoons L : \gamma \qquad \mathcal{K}_3.$$

The model is described by the following polynomial:

$$0 = -N_\gamma + [\gamma] + \mathcal{K}_3[L][\gamma]. \tag{19}$$

The signaling complex is characterized by the quantity $\mathcal{K}_3[L][\gamma]$.

***F.2. Homo-dimeric RTK.*** In the homo-dimeric RTK model, the primary chain, $\gamma$, dimerizes before binding to the ligand:

$$\begin{aligned} \gamma + \gamma &\rightleftharpoons \gamma : \gamma && \mathcal{K}_2, \\ L + \gamma : \gamma &\rightleftharpoons \gamma : \gamma : L && \mathcal{K}_3. \end{aligned}$$

The model is described by the following polynomial:

$$0 = -N_\gamma + [\gamma] + 2\mathcal{K}_2(\mathcal{K}_3[L] + 1)[\gamma]^2. \tag{20}$$

The number of signaling complexes is characterized by the quantity $\mathcal{K}_2\mathcal{K}_3[L][\gamma]^2$.

***F.3. Hetero-dimeric RTK.*** We called "hetero-dimeric RTK" the model of the receptor which associates a $\gamma$ and an $\alpha$ chain, and has intrinsic kinase activity. The biochemical reactions with their affinity constants are:

$$\begin{aligned} \gamma + \alpha &\rightleftharpoons \gamma : \alpha && \mathcal{K}_2, \\ L + \gamma : \alpha &\rightleftharpoons \gamma : \alpha : L && \mathcal{K}_3. \end{aligned}$$

The model is described by the following polynomial system:

$$\begin{aligned} 0 &= - N_\alpha + [\alpha] + \mathcal{K}_2[\gamma][\alpha] + \mathcal{K}_3\mathcal{K}_2[L][\gamma][\alpha], \\ 0 &= - N_\gamma + [\gamma] + \mathcal{K}_2[\gamma][\alpha] + \mathcal{K}_2\mathcal{K}_3[L][\gamma][\alpha]. \end{aligned} \tag{21}$$

The number of signaling complexes is characterized by the quantity $K_2K_3[L][\gamma][\alpha]$.

***F.4. Homo-dimeric receptor with JAK.*** The so-called "homo-dimeric JAK" model describes a homo-dimeric receptor which requires a downstream kinase bound to the $\gamma$ chain to signal. There is now the possibility to create a signaling complex $L:\gamma:j:\gamma:j$, as well as two different dummy complexes $L:\gamma:\gamma$ and $L:\gamma:\gamma:j$. The biochemical reactions scheme is as follows:

$$\begin{array}{lcll} j+\gamma & \rightleftharpoons & j:\gamma & \mathcal{K}_1, \\ j:\gamma+\gamma & \rightleftharpoons & \gamma:\gamma:j & \mathcal{K}_2, \\ j:\gamma+j:\gamma & \rightleftharpoons & \gamma:j:\gamma:j & \mathcal{K}_2, \\ \gamma+\gamma & \rightleftharpoons & \gamma:\gamma & \mathcal{K}_2, \\ L+\gamma:\gamma & \rightleftharpoons & L:\gamma:\gamma & \mathcal{K}_3, \\ L+\gamma:\gamma:j & \rightleftharpoons & L:\gamma:\gamma:j & \mathcal{K}_3, \\ L+\gamma:j:\gamma:j & \rightleftharpoons & L:\gamma:j:\gamma:j & \mathcal{K}_3. \end{array}$$

The polynomial system describing the model is:

$$\begin{aligned} 0 =& -N_\gamma + [\gamma] + \mathcal{K}_1[\gamma][j] + 2\left(\mathcal{K}_1\mathcal{K}_2[\gamma]^2[j] + \mathcal{K}_2\mathcal{K}_1^2[\gamma]^2[j]^2 + K_2[\gamma]^2 + \mathcal{K}_3\mathcal{K}_2[L][\gamma]^2 + \mathcal{K}_3\mathcal{K}_2\mathcal{K}_1[L][\gamma]^2[j] + \mathcal{K}_3\mathcal{K}_2\mathcal{K}_1^2[L][\gamma]^2[j]^2\right), \\ 0 =& -N_j + [j] + \mathcal{K}_1[\gamma]j + K_1K_2[\gamma]^2[j] + 2\mathcal{K}_1^2\mathcal{K}_2[\gamma]^2[j]^2 + \mathcal{K}_3\mathcal{K}_2\mathcal{K}_1[L][\gamma]^2[j] + 2\mathcal{K}_3\mathcal{K}_2\mathcal{K}_1^2[L][\gamma]^2[j]^2. \end{aligned} \quad [22]$$

The number of signaling complexes is characterized by the quantity $\mathcal{K}_3\mathcal{K}_2\mathcal{K}_1^2[L][\gamma]^2[j]^2$.

***F.5. Hetero-dimeric receptor with JAK.*** The hetero-dimeric JAK model is similar to model I: we renamed the species and the affinity constants to match the other models. The biochemical reaction scheme is as follows:

$$\begin{array}{lcll} j+\gamma & \rightleftharpoons & j:\gamma & \mathcal{K}_1, \\ j:\gamma+\alpha & \rightleftharpoons & \alpha:\gamma:j & \mathcal{K}_2, \\ \gamma+\alpha & \rightleftharpoons & \alpha:\gamma & \mathcal{K}_2, \\ L+\alpha:\gamma & \rightleftharpoons & L:\alpha:\gamma & \mathcal{K}_3, \\ L+\alpha:\gamma:j & \rightleftharpoons & L:\alpha:\gamma:j & \mathcal{K}_3. \end{array} \quad [23]$$

The model is described by the following polynomial system:

$$\begin{aligned} 0 &= -N_\gamma + [\gamma] + \mathcal{K}_2[\gamma][\alpha] + \mathcal{K}_1[\gamma][j] + \mathcal{K}_2\mathcal{K}_1[\gamma][\alpha][j] + \mathcal{K}_3\mathcal{K}_2 L[\gamma][\alpha] + \mathcal{K}_3\mathcal{K}_2\mathcal{K}_1[L][\gamma][\alpha][j], \\ 0 &= -N_\alpha + [\alpha] + \mathcal{K}_2[\gamma][\alpha] + \mathcal{K}_2\mathcal{K}_1[\gamma][\alpha][j] + \mathcal{K}_3\mathcal{K}_2[L][\gamma][\alpha] + \mathcal{K}_3\mathcal{K}_2\mathcal{K}_1[L][\gamma][\alpha][j], \\ 0 &= -N_j + [j] + \mathcal{K}_1[\gamma][j] + \mathcal{K}_2\mathcal{K}_1[\gamma][\alpha][j] + \mathcal{K}_3\mathcal{K}_2\mathcal{K}_1[L][\gamma][\alpha][j]. \end{aligned} \quad [24]$$

The quantity $\mathcal{K}_3\mathcal{K}_2\mathcal{K}_1[L][\gamma][\alpha][j]$ characterizes the number of signalling complexes.

**G. Parameter sensitivity analysis.** To assess the influence of the model parameters on the amplitude and $\text{EC}_{50}$, we performed a global stability analysis for each model presented in the paper (except for the allostery model). Of particular interest is the relative importance of the total number of the receptor subunit chains and the kinase ($N_i$), where modelled. Affinity constants

($K_i$) are, in principle, fixed, measured values; however, we included them in our analysis to gain information about the effects of fluctuations and measurement errors. We compared three methods of global stability analysis, the classic Sobol indices (5), the eFAST algorithm (6), and an indirect method via a lightGBM model (7). To implement the Sobol and eFAST methods we used the Python package `SALib` (8). For each method we created data points sampled in parameter space from the hypercube defined by Table S2. We then solved the model equations numerically to obtain amplitude and $EC_{50}$ values.

Let us denote by $n_{params}$ the number of parameters in a model. For the `SALib`-based methods, we used the package-internal sampler to sample $1,024 \times (2n_{params} + 2)$ sets for the Sobol method and $1.1 \times 10^4 \times n_{params}$ for the eFAST method. These numbers of samples proved sufficient to obtain reliable sensitivity scores. However, the sensitivity analysis proved numerically unstable for some models due to the difficulty of solving the polynomial equations with sufficient computational accuracy. Therefore, we do not show these results here. In the cases where the Sobol and eFAST methods converged, we found good agreement with the lightGBM method.

More control is given in the lightGBM method, where we sample parameter configurations by Latin hypercube sampling. We turned the parameter importance problem into a prediction one, where amplitude and $EC_{50}$ are predicted by two separate machine-learning models (which we call trees here to avoid confusion). An advantage of this method is that with decision-tree-based methods, we get two pieces of information. We get a prediction score, to evaluate the accuracy of the model. Feature importances come for free with the fitted trees. We use the prediction accuracy as a proxy for reliability of the sensitivity analysis. We first created a data set of $2 \times 10^5$ points, and we also encountered the issue of numerical instabilities when solving the polynomial equations, in particular, when the $EC_{50}$ values were found to be comparatively large. We circumvented this problem by pruning large $EC_{50}$ values.

We first determined the pruning thresholds. We fitted the trees $10^2$ times for each model and in each fit, we created a training set by sampling $10^5$ data points from our data set, and an independent validation set of $2 \times 10^4$ data points. We determined the threshold for pruning by evaluating the predictive performance of the trees and required a minimum average variance explained ($R^2$) of 0.85 for each individual model. This resulted in removing all data points with an $EC_{50} > 10^{-9}$ in model I and model II.

We repeated the experiments on the final data set and obtained the parameter importances (where, again, we fitted each tree $10^2$ times to assess the stability of the method). In general, the prediction accuracy was satisfactory with a variance explained of mostly 0.99. Model I and model II are an exception with a lowest average $R^2$ of 0.87 on the $EC_{50}$ (data not shown). We then normalised the parameter importances to one in order to obtain the relative parameter importances. The results of these analyses are plotted in Fig. S10.

One aspect to note is that not only is the numerical stability greatly enhanced when analytical expressions for the amplitude and $EC_{50}$ were obtained, but the data generation process is also much faster. However, analytic expressions have not been obtained for all models (in particular, the homo-dimeric RTK model) and, therefore, for consistency and comparability, we use only the numerical solutions.

| | $N_\gamma$ | $N_\alpha$ | $N_j$ | $N_x$ | $K_1$ | $K_2$ | $K_3$ | $K_4$ | $K_5$ | $K_6$ | $K_7$ |
|---|---|---|---|---|---|---|---|---|---|---|---|
| **Lower bound** | 10 | 10 | 10 | 10 | $10^{-6}$ | $10^{-6}$ | $10^{-6}$ (with x),<br>$10^{9}$ (otherwise) | $10^{4}$ | $10^{4}$ | $10^{-4}$ | $10^{-4}$ |
| **Upper bound** | $10^{6}$ | $10^{6}$ | $10^{6}$ | $10^{6}$ | $10^{-2}$ | $10^{-2}$ | $10^{-2}$ (with x),<br>$10^{11}$ (otherwise) | $10^{11}$ | $10^{11}$ | 10 | $10^{-1}$ |

**Table S2. The hypercube of parameters.**

**H. Summary of the models: amplitude and EC$_{50}$ values.** Table S3 summarizes the biochemical reaction schemes of the models studied in this paper. Table S4 recapitulates the amplitude and EC$_{50}$ analytic expressions obtained for the different models when it was feasible. Expressions for homo-dimeric and hetero-dimeric RTK models were obtained following our method described in Ref. (9). The computation of the amplitude and EC$_{rm50}$ expressions for model I and model II were obtained in Ref. (9).

| Model | Equations | | | |
|---|---|---|---|---|
| Monomeric | $L + \gamma : \gamma$ | $\rightleftharpoons$ | $L : \gamma : \gamma$ | $\mathcal{K}_3$ |
| Homo-dimeric RTK | $\gamma + \gamma$ | $\rightleftharpoons$ | $\gamma : \gamma$ | $\mathcal{K}_2$ |
| | $L + \gamma : \gamma$ | $\rightleftharpoons$ | $L : \gamma : \gamma$ | $\mathcal{K}_3$ |
| Hetero-dimeric RTK | $\gamma + \alpha$ | $\rightleftharpoons$ | $\gamma : \alpha$ | $\mathcal{K}_2$ |
| | $L + \gamma : \alpha$ | $\rightleftharpoons$ | $L : \gamma : \alpha$ | $\mathcal{K}_3$ |
| Homo-dimeric JAK | $j + \gamma$ | $\rightleftharpoons$ | $j : \gamma$ | $\mathcal{K}_1$ |
| | $j : \gamma + \gamma$ | $\rightleftharpoons$ | $\gamma : \gamma : j$ | $\mathcal{K}_2$ |
| | $j : \gamma + j : \gamma$ | $\rightleftharpoons$ | $j : \gamma : \gamma : j$ | $\mathcal{K}_2$ |
| | $\gamma + \gamma$ | $\rightleftharpoons$ | $\gamma : \gamma$ | $\mathcal{K}_2$ |
| | $L + \gamma : \gamma$ | $\rightleftharpoons$ | $L : \gamma : \gamma$ | $\mathcal{K}_3$ |
| | $L + \gamma : \gamma : j$ | $\rightleftharpoons$ | $L : \gamma : \gamma : j$ | $\mathcal{K}_3$ |
| | $L + \gamma : j : \gamma : j$ | $\rightleftharpoons$ | $L : \gamma : j : \gamma : j$ | $\mathcal{K}_3$ |
| Hetero-dimeric JAK (model I) | $j + \gamma$ | $\rightleftharpoons$ | $j : \gamma$ | $\mathcal{K}_1 = K_1$ |
| | $j : \gamma + \alpha$ | $\rightleftharpoons$ | $\alpha : \gamma : j$ | $\mathcal{K}_2 = K_2$ |
| | $\gamma + \alpha$ | $\rightleftharpoons$ | $\alpha : \gamma$ | $\mathcal{K}_2 = K_3 = K_2$ |
| | $L + \alpha : \gamma$ | $\rightleftharpoons$ | $L : \alpha : \gamma$ | $\mathcal{K}_3 = K_4$ |
| | $L + \alpha : \gamma : j$ | $\rightleftharpoons$ | $L : \alpha : \gamma : j$ | $\mathcal{K}_3 = K_5 = K_4$ |
| Allostery model | $j_3 + \gamma_c$ | $\rightleftharpoons$ | $j_3 : \gamma_c$ | $K_1$ |
| | $j_3 : \gamma_c + \alpha_7$ | $\rightleftharpoons$ | $\alpha_7 : \gamma_c : j$ | $K_2$ |
| | $\gamma_c + \alpha_7$ | $\rightleftharpoons$ | $\alpha_7 : \gamma_c$ | $K_3$ |
| | $L + \alpha_7 : \gamma_c$ | $\rightleftharpoons$ | $L : \alpha_7 : \gamma_c$ | $K_4$ |
| | $L + \alpha_7 : \gamma_c : j_3$ | $\rightleftharpoons$ | $L : \alpha_7 : \gamma_c : j_3$ | $K_5$ |
| Model II | $j_3 + \gamma_c$ | $\rightleftharpoons$ | $j_3 : \gamma_c$ | $K_1$ |
| | $j_3 : \gamma_c + \alpha_7$ | $\rightleftharpoons$ | $\alpha_7 : \gamma_c : j_3$ | $K_2$ |
| | $\gamma_c + \alpha_7$ | $\rightleftharpoons$ | $\alpha_7 : \gamma_c$ | $K_3 = K_2$ |
| | $L + \alpha_7 : \gamma_c$ | $\rightleftharpoons$ | $L : \alpha_7 : \gamma_c$ | $K_4$ |
| | $L + \alpha_7 : \gamma_c : j_3$ | $\rightleftharpoons$ | $L : \alpha_7 : \gamma_c : j$ | $K_5 = K_4$ |
| | $x + \gamma_c$ | $\rightleftharpoons$ | $x : \gamma_c$ | $K_6$ |
| | $x + \gamma_c : j_3$ | $\rightleftharpoons$ | $x : \gamma_c : j_3$ | $K_7 = K_6$ |

**Table S3. Biochemical reaction scheme of the receptor models considered in the paper.**

| **Model** | **Amplitude** | **$\mathbf{EC}_{50}$** |
|---|---|---|
| Monomeric | $N_\gamma$ | $\frac{1}{\mathcal{K}_3}$ |
| Homo-dimeric RTK | $\frac{N_\gamma}{2}$ | $\frac{1+2\mathcal{K}_2 N_\gamma+\sqrt{1+4\mathcal{K}_2 N_\gamma}}{2\mathcal{K}_2\mathcal{K}_3 N_\gamma}$ |
| Hetero-dimeric RTK | $M$ | $M\frac{1+\mathcal{K}_2(N_\gamma+N_\alpha-M)+\sqrt{1+\mathcal{K}_2^2(N_\alpha-N_\gamma)^2+2\mathcal{K}_2(N_\gamma+N_\alpha-M)}}{\mathcal{K}_2\mathcal{K}_3(M-2N_\gamma)(M-2N_\alpha)}$ |
| Homo-dimeric JAK | NS | NS |
| Hetero-dimeric JAK (model I) | $\frac{K_1 z}{1+K_1 z}M$ | $M\frac{1+\mathcal{K}_2(N_\gamma+N_\alpha-M)+\sqrt{1+\mathcal{K}_2^2(N_\alpha-N_\gamma)^2+2\mathcal{K}_2(N_\gamma+N_\alpha-M)}}{\mathcal{K}_2\mathcal{K}_3(M-2N_\gamma)(M-2N_\alpha)}$ |
| Allostery model | NS | NS |
| Model II | $\frac{K_1 z}{1+K_1 z}M$ | Problem of finding $EC_{50}$ value reduced to solving a polynomial of degree $3$ |

**Table S4. Amplitude and $EC_{50}$ values for the receptor models considered in this paper. We have: $z = \frac{-1+K_1(N_j-N_\gamma)+\sqrt{\Delta}}{2K_1}$, $\Delta = 4K_1N_j + (1+K_1(N_\gamma - N_j))^2$, and $M = \min(N_\gamma, N_\alpha)$. The notation used here for the affinity constants is the same as in Table S3 ($K_1 = \mathcal{K}_1$). Expressions of the amplitude and $EC_{50}$ for model I and model II were obtained in Ref. (9). The other expressions were computed following the method described in the same reference. Model homo-dimeric JAK and the allostery model lead to Gröbner bases involving high degree polynomials and thus, analytic expressions of the amplitude and $EC_{50}$ were not found (NS=not solvable).**

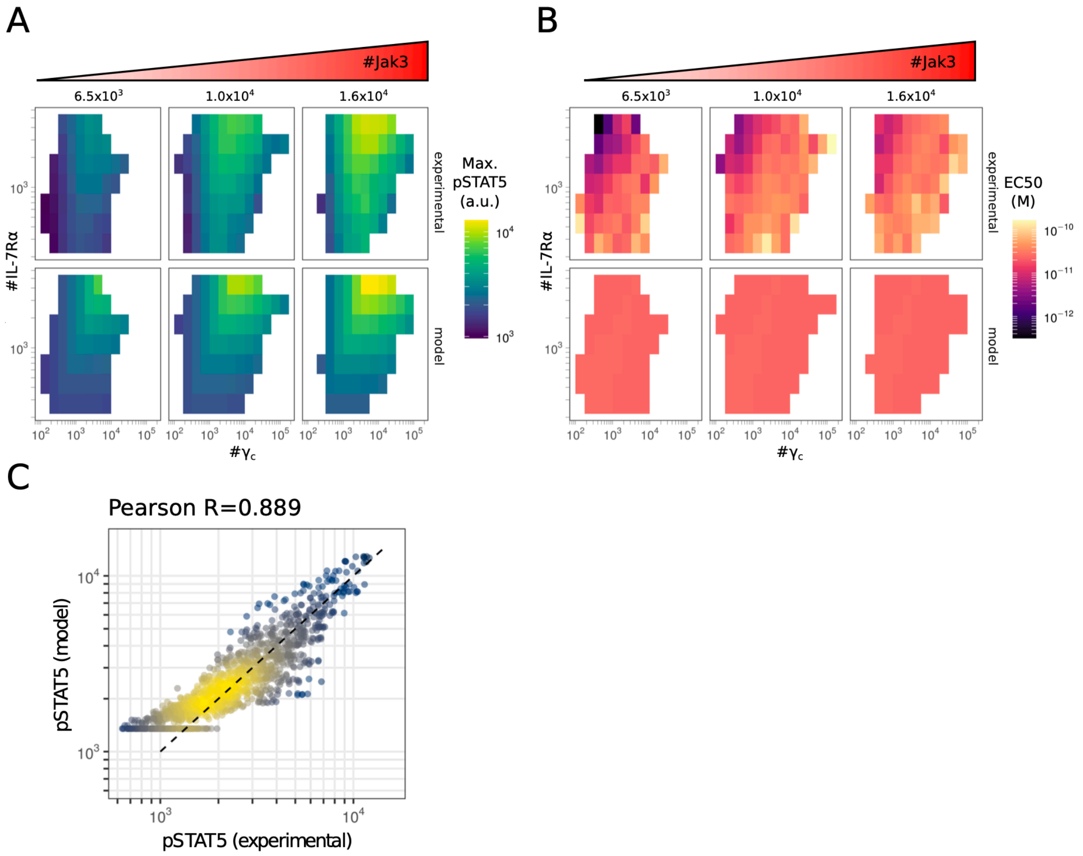


**Fig. S7.** A, B – Heatmaps showing both experimental and model estimates of the maximum pSTAT5 intensity (A) and IL-7 EC$_{50}$ (B) as a function of the number of $\gamma_c$, IL-7R$\alpha$ and JAK3 molecules per cell. Each bin corresponds to a subpopulation of cells whose number of $\gamma_c$, IL-7R$\alpha$ and JAK3 molecules per cell lie within a given distance from the bin center (see methods) as indicated on top for JAK3, and on the $x$ and $y$ axis for IL-7R$\alpha$ and $\gamma_c$, respectively. For each bin, experimental values of the maximum pSTAT5 intensity and of the IL-7 EC$_{50}$ are estimated from a fit of the experimental pSTAT5 to a non-cooperative Hill function (see Methods). Model estimates of the maximum pSTAT5 intensity and of the IL-7 EC$_{50}$ are computed from the theoretical pSTAT5 values corresponding to the best fit of the first model with all parameters fitted with no priors (see Methods for more details on model definition and the entry "model_I_full" in file Supp_table_1 for a summary of model parameters and fit scores). C – Comparison between experimental and modeled pSTAT5 intensities across all bins and all conditions of stimulation ($n = 1,372$ points). Modeled pSTAT5 intensities correspond to the same model as in A and B.

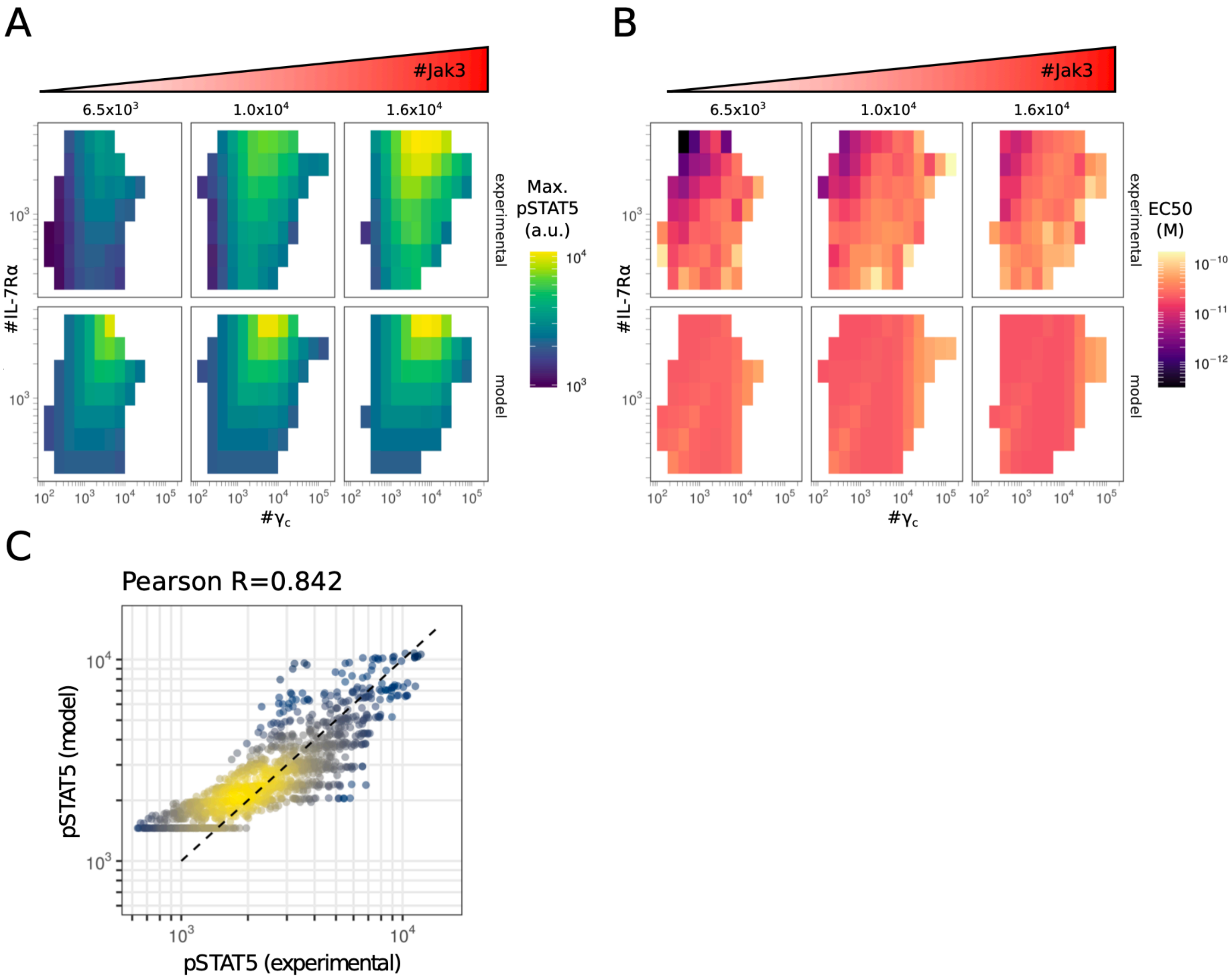


**Fig. S8.** A, B – Heat maps showing both experimental and model estimates of the maximum pSTAT5 intensity (A) and IL-7 $EC_{50}$ (B) as a function of the number of $\gamma_c$, IL-7R$\alpha$ and JAK3 molecules per cell. Each bin corresponds to a subpopulation of cells whose number of $\gamma_c$, IL-7R$\alpha$ and JAK3 molecules per cell lie within a given distance from the bin center (see methods), as indicated on top for JAK3, and on the $x$ and $y$ axis for IL-7R$\alpha$ and $\gamma_c$, respectively. For each bin, experimental values of the maximum pSTAT5 intensity and of the IL-7 $EC_{50}$ are estimated from a fit of the experimental pSTAT5 to a non-cooperative Hill function (see Methods). Model estimates of the maximum pSTAT5 intensity and of the IL-7 $EC_{50}$ are computed from the theoretical pSTAT5 values corresponding to the best fit of the allostery model with all model parameters being fitted with no priors (see Methods for more details on model definition and the entry "model_I_allo_full" in file Supp_table_1 for a summary of model parameters and fit scores). C – Comparison between experimental and modeled pSTAT5 intensities across all bins and all conditions of stimulation ($n = 1,372$ points). Modeled pSTAT5 intensities correspond to the unconstrained fit of the allostery model with all parameters as in A and B. The corresponding Pearson R correlation coefficient is also indicated.

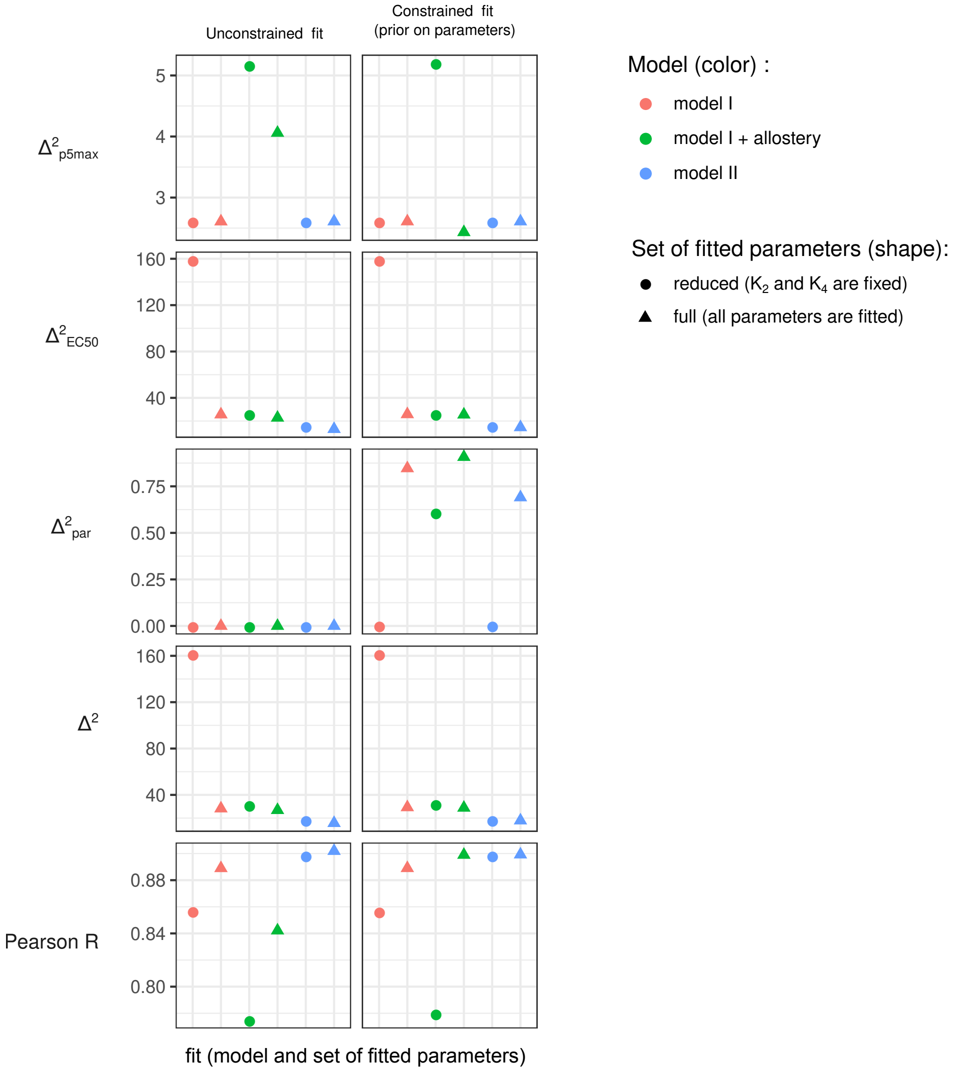


**Fig. S9.** Results of the different fits. Scores assessing fit quality are represented for the different fits performed in this study (values can be found in Supplementary file Supp_table_1). See Methods for the definition of the sum of squared distances $\Delta^2_{p5_{max}}$, $\Delta^2_{EC_{50}}$, $\Delta^2_{par}$ and $\Delta^2$. The Pearson $R$ coefficient corresponds to the comparison between log10-transformed experimental and model estimates of pSTAT5 intensities across all bins and conditions of stimulation.

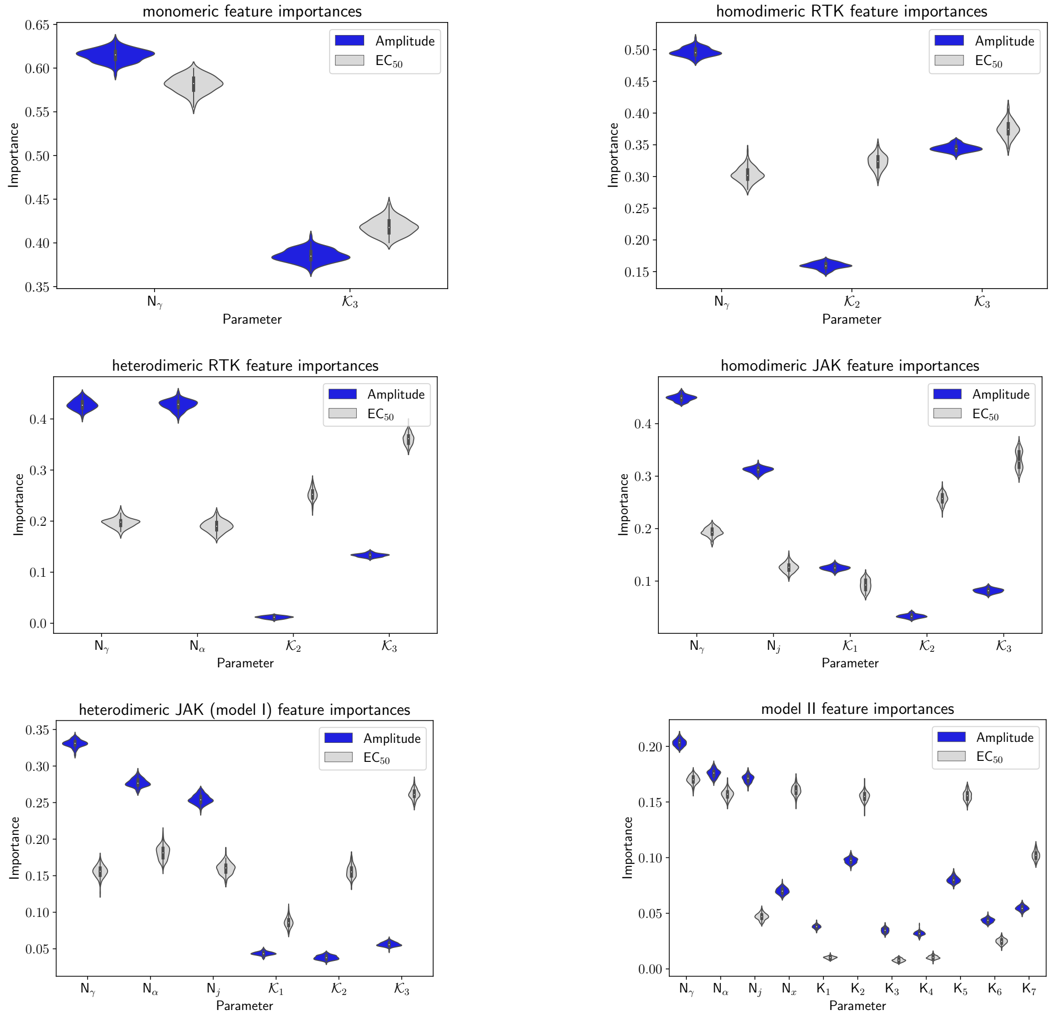


**Fig. S10.** The feature importances of the various models (parameter notation taken from Table S3). We observe that the individual affinity constants only play a minor role in the determination of amplitude, however, they tend to have more importance in the $EC_{50}$ value.
